# DNA methyltransferase 1 (DNMT1) function is implicated in the age-related loss of cortical interneurons

**DOI:** 10.1101/2020.03.06.981290

**Authors:** Anne Hahn, Cathrin Bayer, Daniel Pensold, Jessica Tittelmeier, Lisa Marx-Blümel, Lourdes González-Bermúdez, Jenice Linde, Jonas Groß, Gabriela Salinas-Riester, Thomas Lingner, Julia von Maltzahn, Marc Spehr, Tomas Pieler, Anja Urbach, Geraldine Zimmer-Bensch

## Abstract

Increased life expectancy in modern society comes at the cost of age-associated disabilities and diseases. Aged brains not only show reduced excitability and plasticity, but also a decline in inhibition. Age-associated defects in inhibitory circuits likely contribute to cognitive decline and age-related disorders. Molecular mechanisms that exert epigenetic control of gene expression, contribute to age-associated neuronal impairments. Both DNA methylation, mediated by DNA methyltransferases (DNMTs), and histone modifications maintain neuronal function throughout lifespan. Here we provide evidence that DNMT1 function is implicated in the age-related loss of cortical inhibitory interneurons. Deletion of *Dnmt1* in parvalbumin-positive interneurons attenuates their age-related decline in the cerebral cortex. Moreover, DNMT1-deficient mice show improved somatomotor performance and reduced aging-associated transcriptional changes. A decline in the proteostasis network, responsible for the proper degradation and removal of defective proteins, is suggested to be essentially implicated in age- and disease-related neurodegeneration. Our data suggest that DNMT1 acts indirectly on interneuron survival in aged mice by modulating the proteostasis network during life-time.

## 1 Introduction

Aging mediates structural, neurochemical and physiological alterations in the brain that are associated with behavioral changes, memory decline, and cognitive impairments (Rozycka & Liguz-Lecznar, 2017). Cognitive aging results in metabolic, hormonal and immune dysregulation, increased oxidative stress and inflammation, altered neurotransmission and synaptic plasticity as well as reduced neurotrophic support of neurons (Rozycka & Liguz-Lecznar, 2017). Notably, in the aging brain, distinct cell types and circuits are affected differently (reviewed in Zimmer-Bensch, 2019).

Inhibitory interneurons of the cerebral cortex are key players in information processing (Kann et al., 2014) and particularly affected by aging. Reduced interneuron numbers were reported across diverse species and cortical regions (reviewed in Zimmer-Bensch, 2018). Additionally, morphological abnormalities and synaptic dysfunction of GABAergic synapses emerge as major factors in aging-related impairments of nervous system function (Morrison & Baxter, 2012). These findings are in agreement with the decline in inhibition reported by several studies (Cheng & Lin, 2013; Shetty & Turner, 1998; Stanley & Shetty, 2004). In line with reduced neurotransmitter release, major changes in the expression of genes related to neurotransmission and transcriptional repression of GABA-related transcripts have been described for the human prefrontal cortex (Loerch et al., 2008), but also in brains across different mammalian species (reviewed in Zimmer-Bensch, 2019). Diminished expression of genes involved in synaptic function indeed appears to be a conserved feature of mammalian brain aging (Ianov et al., 2016; Jiang et al., 2001; Loerch et al., 2008).

Given the importance of GABAergic inhibitory interneurons in cortical information processing, age-associated defects in inhibitory circuits contribute to cognitive decline and age-related disorders (Rozycka & Liguz-Lecznar, 2017). Such defects include the loss of synaptic contacts, decreased GABA release, and reduced postsynaptic responsiveness, thus disturbing the excitation/inhibition balance in the aging brain. Fast-spiking parvalbumin (PV) positive interneurons represent the most abundant subset of cortical inhibitory interneurons (Druga, 2009). They execute both feedforward and feedback inhibition, and are responsible for generating gamma-frequency oscillations (Buzsáki & Wang, 2012; Kann et al., 2014; Sohal et al., 2009; Willems et al., 2018). In schizophrenia patients, a reduction in PV interneurons and their dysfunction have been associated with the loss of gamma oscillations, manifesting in working memory and executive function deficits (Sohal et al., 2009; Torrey et al., 2005). Upon aging, PV interneurons are diminished in cell numbers in the somatosensory, auditory, and motor cortices of rats as well as in the hippocampus (Miettinen et al., 1993; Ouda et al., 2008). Moreover, altered PV interneuron function is implicated in age-related diseases like Alzheimer’s Disease (AD; Rossignol, 2011; Verret et al., 2012). Together, these studies emphasize the role PV interneurons have in cortical function. Hence, detailed analysis of age-related changes in the PV interneuron subpopulation might help to understand the processes underlying cognitive aging and age-related memory impairments.

Apart from synaptic defects, aging is accompanied by a declining proteostasis network that causes ineffective protein degradation, which can lead to neuronal death (Douglas & Dillin, 2010). Lysosomal degradation is of great importance for removing defective proteins or protein aggregates delivered by autophagy- or endocytosis-triggered endosomal pathways (Nixon et al., 2000; Nixon & Cataldo, 1995; Winckler et al., 2018), and lysosomal dysfunction is associated with age-related neurodegenerative pathologies like Parkinson’s and Alzheimer’s disease (Carmona-Gutierrez et al., 2016; Zhang et al., 2009). Besides lysosomal degradation, protein removal can be achieved by inclusion into multivesicular bodies (MVBs), which are then released as exosomes into the extracellular space (Riva et al., 2019). Exosomal release has recently been implicated in contributing to neurodegenerative disease and mental disorders (Bellingham et al., 2012; Delpech et al., 2019; Saeedi et al., 2019).

At the molecular level, epigenetic mechanisms emerge as crucial players in the physiology of healthy aging as well as in the pathophysiology of age-related neurological disorders. Epigenetic mechanisms involve inheritable as well as reversible chromatin modifications, including DNA methylation and histone modifications, which influence gene transcription and post-transcriptional events (Fuks, 2005)(**REFS**). Further epigenetic key players are represented by non-coding RNAs, which can act on transcriptional, post-transcriptional and translational level (Cech & Steitz, 2014; Geisler & Coller, 2013; Zimmer-Bensch, 2019).

DNA-methylation executed by DNMTs affects gene expression through diverse mechanisms (Gelfman et al., 2013; Lyko, 2018; Maunakea et al., 2010) and is implicated in the pathogenesis of brain aging (Cui & Xu, 2018). We have recently found that DNA methyltransferase 1 (DNMT1)-dependent DNA methylation modulates synaptic function of cortical PV interneurons by acting on endocytosis-mediated vesicle recycling (Pensold et al., n.d.). Since synaptic function regulation and DNA methylation are involved in brain aging, we here investigated whether DNMT1-dependent gene regulation in PV interneurons contributes to their age-related defects.

## 2 Methods

### Animals

The following mouse strains were used: C57BL/6 wild-type mice and transgenic mice on the C57BL/6 background including *Pvalb-Cre*/*tdTomato*/*Dnmt1* control as well as *Pvalb-Cre*/*tdTomato*/*Dnmt1 loxP*^*2*^ mice. The transgenic mice were established by crossing the *Pvalb-Cre* line (obtained from Christian Huebner, University Hospital Jena, Germany and described in Hippenmeyer et al., 2005) with the *tdTomato* transgenic reporter mice (obtained from Christian Huebner, University Hospital Jena, Germany and described in Madisen et al., 2010) and the *Dnmt1 loxP*^*2*^ mice, (B6;129Sv-Dnmt1^tm4Jae^/J, Jaenisch laboratory, Whitehead Institute; USA). The *Dnmt1 loxP*^*2*^ mice have LoxP-sites flanking exons 4 and 5 of the *Dnmt1* gene. CRE-mediated deletion leads to out-of-frame splicing from exon 3 to exon 6, resulting in a null *Dnmt1* allele (Jackson-Grusby et al., 2001). Transgenic *Pvalb-Cre*/*tdTomato*/*Dnmt1* control and *Pvalb-Cre*/*tdTomato*/*Dnmt1 loxP*^*2*^ mice are abbreviated as *Dnmt1* WT (*wild-type*) and *Dnmt1* KO (*knockout*) in the figures, respectively. The floxed *Dnmt1* allele was genotyped with forward GGGCCAGTTGTGTGACTTGG and reverse CCTGGGCCTGGATCTTGGGGA primer pairs resulting in a 334 bp WT and 368 bp mutant band. The *tdTomato* allele was genotyped using the following set of four primers: WT forward AAGGGAGCTGCAGTGGAGTA, WT reverse CCGAAAATCTGTGGGAAGTC, mutant forward CTGTTCCTGTACGGCATGG, mutant reverse CTGTTCCTGTACGGCATGG giving WT (297 bp) and mutant (196 bp) bands. The *Pvalb*-*Cre* genotyping was performed by applying AAACGTTGATGCCGGTGAACGTGC forward and TAACATTCTCCCACCGTCAGTACG reverse primer resulting in a 214 bp fragment. All animal procedures were performed in strict compliance with the EU directives 86/609/EWG and 2007/526/EG guidelines for animal experiments and were approved by the local government (Thueringer Landesamt, Bad Langensalza, Germany). Animals were housed under 12 h light/dark conditions with *ad libitum* access to food and water.

### Ladder Rung Test

Cohorts of *Pvalb-Cre/tdTomato/Dnmt1* WT as well as *Pvalb-Cre/tdTomato/Dnmt1 loxP*^*2*^ mice were consecutively tested over different ages starting from 3 months to 21 months. Mice were placed onto a ladder beam (transparent) with rungs in a regular pattern (every 10 mm) at a slight incline (∼30°) with the home box at the end. Time to cross the ladder was measured, not including the time spent in a stop or walking back towards the starting point. The scoring system according to Metz & Whishaw (2009) was used for foot placement accuracy. In each test session the animals had to cross the ladder consecutively for three times.

### Isolation and Primary Cultivation of Dissociated Embryonic Single Cells

Pregnant dams were anesthetized by an intraperitoneal injection of 50% chloral hydrate in phosphate buffered saline (PBS; pH 7.4; 2.5 µg chloral hydrate per g body weight). After death of the dam, all embryos were dissected out of both uterine horns and instantly decapitated. The brain was dissected in ice-cold and sterile filtered Gey’s Balanced Salt Solution (GBSS; 1.53 mM CaCl_2_, 3.66 mM KCl, 0.22 mM KH_2_PO_4_, 1.03 mM MgCl_2_ × 6H_2_O, 0.28 mM MgSO_4_ × 7H_2_O, 137.93 mM NaCl, 2.702 mM NaHCO_3_, 0.84mM Na_2_HPO_4_, and 5.56 mM D(+)-Glucose).

Dissociated embryonic medial ganglionic eminence (MGE)-derived single cells for primary culture were prepared from MGE explants dissected from coronal brain sections according to (Zimmer et al., 2011). Briefly, embryonic brains were prepared in Krebs buffer (126 mM NaCl, 2.5 mM KCl, 1.2 mM NaH_2_PO_4_, 1.2 mM MgCl_2_, 2.1 mM CaCl_2_, 10 mM D(+)-Glucose, and 12.5 mM NaHCO_3_), embedded in 4% low-melt agarose (Carl Roth, Germany) at 37 °C for coronal sectioning with a vibratome at 4 °C. MGE explants were collected in ice-cold Hanks Balanced Salt Solution (HBSS, Invitrogen, USA) supplemented with 0.65% D(+)-Glucose. After incubation with 0.04% trypsin (Invitrogen) in HBSS for 17 min at 37 °C, cells were dissociated by trituration and filtering through nylon gauze (pore size 140 μm; Millipore).

Dissociated neurons were plated on coverslips coated with 19 μg/mL laminin (Sigma-Aldrich, Germany) and 5 μg/mL poly-L-lysine (Sigma-Aldrich) at a density of 225 cells/mm^2^ in Neurobasal Medium (Thermo Fisher Scientific) supplemented with 1xB27 (Thermo Fisher Scientific), 100 U/mL penicillin, 100 μg/mL streptomycin, and 0.5 mM GlutaMax (Thermo Fisher Scientific). After incubation at 37 °C, 5% CO_2_ in a humid atmosphere to 95% H_2_O for 7 days in vitro (DIV), cells were fixed in 4% PFA in PBS (pH 7.4) for 10 min at room temperature.

### Cell culture

Cerebellar granule (CB) cells were cultured in Dulbecco’s Modified Eagle’s Medium with high glucose (DMEM, Invitrogen) supplemented with 10% fetal bovine serum (FBS, Invitrogen), 1% GlutaMAX, 24 mM of KCl, 100 U/mL penicillin, 100 µg/mL streptomycin incubated at 33 °C, 95% H_2_O, 5% CO_2_.

### Transfection with siRNA Oligos and CD63-pEGFP

For siRNA transfections of dissociated embryonic MGE cells of C57BL/6 WT mice and cerebellar granular (CB) cells, reverse lipofection with Lipofectamin© 2000 (Thermo Fisher Scientific, USA) according to the manufacturer’s protocol and as described in (Zimmer et al., 2011) was applied using 15 nM control siRNA (BLOCK-iT Alexa Fluor red or green fluorescent oligo, Invitrogen, USA) and 30 nM *Dnmt1* siRNA, *Rab7* siRNA (Santa Cruz Biotechnology) for 5 h in Opti-MEM I Reduced Serum Medium without antibiotics (Thermo Fisher Scientific). MGE-derived neurons were transfected after 6 DIV, whereas CB cells were plated on coverslips one day prior to transfection. Cells were cultured overnight at 37°C, 95% H_2_O and 5% CO_2_ using the aforementioned cell line specific culture medium prior to fixation.

Transfection for the CD63 overexpression construct was done as described above for siRNA transfection using 2 µg/mL of CD63-pEGFP (Addgene, USA) added for 5 h in Opti-MEM I Reduced Serum Medium (Thermo Fisher Scientific). Cells were cultured overnight at 37 °C, 95% H_2_O and 5% CO_2_ using the aforementioned cell line specific culture medium applied to live cell imaging in a petri dish inserted in a chamber heated to 37 °C using imaging media of Hank’s Balanced Salt Solution (HBSS; Thermo Fisher Scientific) supplemented with 0.65% D(+)-Glucose, 10% FBS, 1% GlutaMAX (Thermo Fisher Scientific), 100 U/mL penicillin, 100 µg/mL streptomycin and 25 µM HEPES (Thermo Fisher Scientific).

### EGF Endocytosis

EGF coupled to Alexa-488 (Molecular Probes, Invitrogen, USA) was used as an endocytic probe. siRNA transfected CB cells were incubated in serum-free DMEM supplemented with 1% BSA for 1 h at 33°C followed by incubation in uptake media (DMEM, 1% BSA, 50 mM HEPES) containing 0.5 µg/mL EGF coupled to Alexa-488 on ice for 1 h. Cells were then washed 3x with ice-cold PBS (pH 7.4) to remove unbound ligands and then incubated for the indicated time points in serum-free DMEM, 1% BSA 1 h at 33°C. Cells were then put on ice, washed 3x ice-cold PBS (pH 7.4), then placed in an acid wash (0.2 M acetic acid, 0.5 M NaCl (pH 2.8)) to remove any non-internalized ligands. After fixation in 4% PFA in PBS (pH 7.4) for 10 min, cells were stained against LAMP1.

### Brain tissue preparation

Mice were deeply anesthetized by intraperitoneal injection of 50% chloral hydrate in phosphate buffered saline (PBS; pH 7.4; 2.5 µg chloral hydrate per g body weight). For *in situ* hybridization experiments, freshly prepared brains were immediately frozen in liquid nitrogen and stored at −80°C. For immunohistochemistry, mice were perfused with PBS (pH 7.4) followed by 4% paraformaldehyde (PFA) in PBS (pH 7.4) and brains were prepared. Post-fixation occurred over night at 4°C. Cryoprotection with 10% and 30% sucrose in PBS overnight was applied before freezing in liquid nitrogen and storage at −80°C.

### *In situ* hybridization, immunohistochemistry and immunocytochemistry

For *in situ* hybridizations, adult brains were cryo-sectioned coronally at −20°C (20 µm). *In situ* hybridizations were performed as described by Zimmer et al. (2011) using digoxigenin-labeled riboprobes. The following primers were used to generate the riboprobe: forward GAGAGCTCTGTCGATGACAGACGTGCTC and reverse GAGGTACCTTCTTCAACCCCAATCTTGC for *Pvalb* (NM_013645.3). The riboprobe was obtained by *in vitro* transcription using DIG-11-UTP (Roche, Germany) from cDNA fragments cloned in pBluescript II SK (Stratagene, USA). For Nissl staining adult brains were cryo-sectioned at −20°C (20 µm) and fixed on slides for 30 min in fixation solution (95% (v/v) ethanol, 5% (v/v) acetic acid). After washing in water, sections were incubated in 0.5% (w/v) cresyl violet for 25 min, and washed in water. Then an ethanol-series (50%, 70% and 99%) was applied for 2.5 min each. Subsequently, sections were incubated in xylol for 5 min and mounted in Depex mounting media (Serva, Germany).

For immunocytochemistry on dissociated MGE cells, permeabilization and washing between different incubation steps was performed with 0.1% (v/v) Triton X-100 in PBS (pH 7.4) for 10 min. Blocking with 5% (v/v) normal goat serum in PBS (pH 7.4) was performed for 30 min and primary antibodies were applied overnight at 4°C, secondary antibodies were applied for 1 h. Cells were washed prior to nuclei staining with DAPI (Molecular Probes, USA) for 5 min. CB cells were permeabilized with 0.2% Triton X-100 in PBS (pH 7.4) for 10 min prior to blocking with 5% normal goat serum in PBS (pH 7.4) for 1 h. Primary antibodies were applied overnight at 4°C, secondary antibodies for 1 h at room temperature (RT). After nuclei staining with DAPI (Molecular Probes, USA) for 5 min, coverslips were embedded in Mowiol (Carl Roth, Germany). Unless noted differently, all steps were performed at room temperature.

The following primary antibodies were used: mouse anti-RFP (1:500, Thermo Fisher Scientific), mouse anti-Parvalbumin (1:2000, Swant Switzerland), rabbit anti-CD63 (1:500, gift from Markus Damme, Biochemisches Institut Christian-Albrechts-Universitaet-Kiel), rat anti-LAMP1 (1:200, Thermo Fisher Scientific).

The following secondary antibodies were applied: goat Alexa 488 anti-mouse (1:1000, Vector), goat Alexa 488 anti-rat (1:1000, Thermo Fisher Scientific), goat Cy3 anti-mouse (1:1000, Jackson Immunoresearch), goat Cy5 anti-mouse (1:1000, Thermo Fisher Scientific), goat Cy5 anti-rabbit (1:1000, Thermo Fisher Scientific).

### Isolation of adult and aged cortical interneurons for FACS

The optimized protocol used to collect the material for DNA and RNA-sequencing was modified based on different protocols (Brewer, 1997; Brewer & Torricelli, 2007; Eide & McMurray, 2005; Saxena et al., 2012). Adult and aged brains were dissected in GBSS (1.53 mM CaCl_2_, 3.66 mM KCl, 0.22 mM KH_2_PO_4_, 1.03 mM MgCl_2_ * 6H_2_O, 0.28 mM MgSO_4_ * 7H_2_O, 137.93 mM NaCl, 2.7 mM NaHCO_3_, 0.84 mM Na_2_HPO_4_, 5.56 mM D(+)-Glucose, pH 7.4) supplemented with 0.65% (D+)-Glucose. Cortical hemispheres were dissected and subsequently handled separately. All following volumes are calculated per cortical hemisphere, which were cut into small pieces and transferred to 5 mL HBSS w/o Ca^2+^ and Mg^2+^ supplemented with 7 mM HEPES, 100 U/mL penicillin, 100 µg/mL streptomycin and 0.65% D(+)-Glucose and washed twice. The tissue was then transferred to 5 mL pre-warmed (20 min at 37°C) Trypsin/EDTA (Life Technologies, USA) supplemented with 132 mM trehalose (Sigma-Aldrich, Germany), 100 U/mL penicillin, 100 µg/mL streptomycin, 10 mM HEPES and 600 U DNase (Applichem, Germany) and incubated for 30 min at 37°C, rotating the samples every 5 min. Samples were washed with 2.1 mL pre-warmed DMEM/F12 supplemented with 10% FBS, 100 U/mL penicillin, 100 µg/mL streptomycin and 132 mM trehalose. After adding 0.9 mL pre-warmed HBSS containing 10 mg/mL Collagenase Type 2 (Worthington, UK) samples were incubated for 25 min at 37°C rotating every 5 min and then washed with 2 mL pre-warmed DMEM/F12 supplemented with 10% FBS, 100 U/mL penicillin, 100 µg/mL streptomycin, 3.3 mM EDTA and 132 mM trehalose prior to cool down on ice for 2 min. Dissolving of samples occurred in 1.5 mL DMEM/F12 supplemented with 10% FBS, 100 U/mL penicillin, 100 µg/mL streptomycin and 132 mM trehalose. Trituration was performed using fire-polished and heat-treated (180°C for 8 h) glass capillaries of three different diameters (about 500 µm, 250 µm and 100 µm), which were coated with DMEM/F12 supplemented with 10% FBS, 100 U/mL penicillin and 100 µg/mL streptomycin prior to use. Mechanical dissociation was performed by pipetting up and down gently 3-5 times for each diameter starting with the largest, avoiding air bubbles. After each step, the supernatant was collected in 1 mL DMEM/F12 supplemented with 10% FBS, 100 U/mL penicillin, 100 µg/mL streptomycin and 132 mM trehalose was added to the original sample. After trituration with the smallest glass capillary, the suspension was filtered through nylon gauze (80-100 µm) and centrifuged for 5 min at 160 g, 4°C. After supernatant removal, the pellet was dissolved in 4 mL HBSS w/o Ca^2+^ and Mg^2+^ supplemented with 7 mM HEPES, 100 U/mL penicillin, 100 µg/mL streptomycin, 0.65% D(+)-Glucose and 132 mM trehalose. After centrifugation (5 min, 160g, 4°C), the pellet was dissolved in PBS (pH 7.4) with 30% Percoll (Sigma-Aldrich, USA) and 132 mM trehalose to perform a density gradient centrifuged for 10 min at 500g and 4°C. The supernatant was removed and the pellet was dissolved in 250 µL HBSS w/o Ca^2+^ and Mg^2+^ supplemented with 7 mM HEPES, 100 U/mL penicillin, 100 µg/mL streptomycin, 0.65% D(+)-Glucose and 132 mM trehalose for FACS.

### FACS enrichment of tdTomato cells

Cell suspensions subjected to FACS were prepared from the cortical hemispheres of adult 6 months and 18 months old *Pvalb-Cre/tdTomato/Dnmt1* WT as well as *Pvalb-Cre/tdTomato/Dnmt1 loxP*^*2*^ mice. Following addition of DAPI, cells were sorted using an ARIA III FACS sorter (BD Biosciences, USA) with a maximal flow rate of 6. The tdTomato reporter was excited by a 561 nm yellow/green solid-state laser and emission signal was detected in a range of 579 nm to 593 nm. According to their forward scatter/side scatter (FCS/SSC) criteria followed by cell doublet exclusion via an FSC-H vs. FSC-W criterium, DAPI-negative living cells were sorted based on a distinctive tdTomato signal. Cells of interest were collected in HBSS w/o Ca^2+^ and Mg^2+^ supplemented with 7 mM HEPES, 100 U/mL penicillin, 100 µg/mL streptomycin, 0.65% D(+)-Glucose and 132 mM trehalose at 4°C and pelleted by centrifugation. Enriched tdTomato cells of one hemisphere were prepared for RNA-sequencing, while cells of the contralateral hemisphere were subjected to DNA-isolation for MeDIP-sequencing for each brain used. For RNA isolation, pellets were dissolved in 500 µL Trizol®Reagent (Life Technologies, USA) and subsequently frozen on dry ice. For MeDIP-Seq analysis, cell pellets were frozen at −80 °C until further use. Only male mice were used for RNA and MeDIP sequencing.

### RNA/DNA isolation of tissue and FAC-sorted cells

Adult cortical hemispheres were dissected from whole brain and frozen in liquid nitrogen as described above. For RNA-sequencing samples were subjected to standard RNA isolation procedure using Trizol®Reagent (Life Technologies, USA). The FACS-enriched tdTomato cells were processed accordingly, with additional application of GlycoBlue (Thermo Fisher Scientific, USA) to a final concentration of 0.2% during RNA precipitation for better visualization of the pellet.

DNA isolation of FACS-enriched tdTomato cells was performed using QIAamp DNA Micro Kit (Quiagen, Germany) according to manufacturer’s instruction and checked for integrity by capillary gel electrophoresis (Bioanalyzer, Agilent Technologies, Inc., USA).

### RNA sequencing of adult cortical tissue

To reveal potentially relevant genes for age related processes in the brain, we performed RNA sequencing of 6 months and 16 months old cortical hemispheres of C57BL/6 mice. The TruSeq RNA Sample Preparation Kit (Illumina, Cat. N°RS-122-2002, USA) was used for library preparation (1 µg total RNA), the QuantiFluor™ dsDNA System (Promega, USA) for quantitation and the DNA 1000 chip on the Bioanalyzer 2100 (Agilent Technologies) to determine the size range of final cDNA libraries prior to amplification and sequencing (cBot and HiSeq2000 from Illumina; PE; 2×100 bp; ca. 30 million reads per sample).

Before alignment of the sequences using Bowtie2; v2.0.2 to the UCSC mouse reference genome mm10, sequences were trimmed for adaptor sequences and phred scores<30 via fastq-mcf (ea-utils v1.1.2-484). Counting the reads to each gene was done via HTSeq python scripts (v0.5.3p9) to the UCSC gene annotation. Data were preprocessed and analyzed in the R/Bioconductor environment (v2.15.2) loading edgeR v.3.0.4 (Robinson et al., 2010).

### RNA Sequencing of FACS-enriched tdTomato cells

RNA was isolated using the Trizol®Reagent protocol according to manufacturer’s instructions. RNA quality was assessed by measuring the RIN (RNA Integrity Number) using the fragment analyzer from Advanced Analytical (USA). Library preparation for RNA-Seq was performed using the TruSeq™ RNA Sample Prep Kit v2 (Illumina, Cat. N°RS-122-2002, USA) starting from 50 ng of total RNA. Accurate quantitation of cDNA libraries was performed by using the QuantiFluor™ dsDNA System (Promega, USA). The size range of final cDNA libraries was determined applying the DNA chip on the fragment analyzer (average 350 bp; Advanced Analytical). cDNA libraries were amplified and sequenced by using the cBot and HiSeq2000 from Illumina (SR; 1×50 bp; ∼30-40 million reads per sample). Sequence images were transformed with Illumina software BaseCaller to bcl files, which were demultiplexed to fastq files with CASAVA v1.8.2. Quality check was done via fastqc (v. 0.10.0, Babraham Bioinformatics, UK). Read alignment was performed using STAR v2.3.0 (Dobin et al., 2013) to the mm10 reference genome. Data were converted and sorted by samtools 0.1.19 and reads per gene were counted via htseq version 0.5.4.p3. Data analysis was performed using R/Bioconductor 3.0.2/2.12 (Luo & Brouwer, 2013); loading DESeq2 (Love et al., 2014). Candidate genes were filtered to an FDR-corrected P value<0.05. Sequence data will be deposited in NCBI’s Gene Expression Omnibus and are accessible through GEO Series upon acceptance of the manuscript.

### MeDIP sequencing of FACS-enriched tdTomato cells

For genome-wide methylation analysis we applied immunoprecipitation methods for the enrichment of 5-methylcytosines. Specifically, 100 ng of genomic DNA were used as starting material. The Methylated-DNA IP Kit from Zymo (Cat. N° D5101) was applied according to manufacturer’s instructions. The product of the IP and control reaction were then used for preparation of Illumina compatible libraries according to the TruSeq Nano DNA Library Prep Kit (Cat. N° FC-121-4001). Libraries were sequenced on a HiSeq 2000 yielding 50 bp single end reads. The sequencing reads were demultiplexed using the Illumina CASAVA tool and sequence quality was checked using FASTQC (http://www.bioinformatics.babraham.ac.uk/projects/fastqc/). The reads were then aligned to the genome of *Mus musculus* (mm10) using Bowtie 2 (versions 2.0.2) with standard parameters. Differentially methylated regions were identified using the MEDIPS package for R version 1.16.0 (Lienhard et al., 2014) with a window size of 700 bp and a minimum coverage of 5% of the window length. A detailed description of the analysis pipeline can be found in Halder et al. (2015).

### Analysis of sequencing data

Normalization of raw counts and differential gene expression analysis were performed using the DESeq2 R package (v 1.12.3; Love et al., 2014). Genes were considered differentially expressed with a Benjamini-Hochberg adjusted P value P<0.05.

Gene list overlaps between differentially expressed and methylated genes took into account that several differentially methylated sites may be annotated to one gene and were quantified using the Jaccard coefficient. Absolute numbers of differentially methylated genes were determined without regard to multiple sites of differential methylation in a single gene. Significance of enrichment of methylated genes was calculated using Fisher’s exact test.

Gene lists were submitted to the *Database for Annotation, Visualization and Integrated Discovery* (DAVID, https://david.ncifcrf.gov) for Gene Ontology or KEGG Pathway term enrichment analysis. Results of GO enrichment analysis were visualized either in a bar diagram including the respective *Benjamini-Hochberg* corrected p-value, the number of genes and the enrichment fold change included in a certain term or using Cytoscape 3.2.1 and the EnrichmentMap plugin (Merico et al., 2010). For visualization, no terms were excluded based on their P value or false discovery rate (FDR). Terms appear as connected if the Jaccard Coefficient of the associated genes was >0.25. The resulting network was manually adapted to exclude non-relevant nodes.

Heat maps were generated using R package pheatmap (https://CRAN.R-project.org/package=pheatmap). For heat maps showing comparison between two datasets, data were normalized to 6 months WT. In case of heat maps illustrating more than two samples, data were scaled. KEGG pathway was visualized using R package pathview (Luo & Brouwer, 2013).

### Microscopy and image data analysis

Images of immunohistochemistry staining of adult tissue sections or immunocytochemistry of stained cell culture was recorded either with an inverted confocal laser scanning microscope TCS SP5 (Leica Microsystems, Germany) or with an inverted transmitted light microscope Axio Cellobserver Z1 equipped with MosaiX module for tile scanning and apotome for confocal like imaging (Carl Zeiss Microscopy, Germany). Photographs were analyzed using the free Fiji software (Schindelin et al., 2012).

For life cell imaging of CB cells transfected with the CD63-pEGFP and either control or Dnmt1 siRNA images were taken with Axio CellObserver Z1, x40 optical magnification using apotome. Z-stack was applied over the whole cell and acquisition was performed every 5 min for 1 h. *.zvi-files were opened with ImageJ; maximum intense projection was performed and data were exported as *.avi with 5 frames per second. The movement of CD63-pEGFP positive vesicles was measured direction specific from one timepoint to the next and speed was calculated based on the time interval.Analysis of cell number in adult sections was performed with ImageJ cell counter plugin. Counted cell numbers in section analysis were normalized to the area of the counted region.

For fluorescence intensity measurements, each experimental design was imaged at one particular microscope with consistent settings regarding exposure time and light intensity at the CellObserver Z1 or laser power, gain and spectral settings at the SP5 LSM. Fluorescence intensity measurement for the CD63 staining and Lamp1 staining was performed in the processes of the cells. For each picture, background correction was performed by subtracting the mean fluorescent intensity from three background areas. Mean fluorescent intensity of the *Dnmt1* siRNA treated cells was normalized to control siRNA. Photoshop CC was applied for image composition. Boxplots were plotted using R. Significance was analyzed with two-tailed Student’s t-test or two-way ANOVA. Significance levels: ∗P < 0.05; ∗∗P < 0.01; and ∗∗∗P < 0.001.

## 3 Results

### Vulnerability of PV-expressing neocortical GABAergic interneurons towards aging

Aging-dependent functional defects in the cortical inhibitory GABAergic system were reported for humans (Cheng & Lin, 2013) as well as for different animal models (Miettinen et al., 1993; Ouda et al., 2008) including mice (Jessen et al., 2017). Since mice serve as key models to study the neurobiology of aging and age-associated neurodegenerative diseases (Bilkei-Gorzo, 2014; Jucker & Ingram, 1997), we tested whether the neocortical GABAergic system is compromised in aged mice. As an initial approach we performed differential gene expression analysis of the whole neocortex from young (6 months) and aged (16 months) C57BL/6 mice. In general, RNA sequencing revealed comparatively low numbers of age-dependent differentially expressed genes (**Fig. 1a**). This was also observed by others when using whole cortical tissue containing a mixed population of cells (e.g. glia versus neurons), which likely show different responses towards aging (Kimmel et al., 2019). Moreover, the aging brain shows mRNA–protein decoupling (Wei et al., 2015), with numerous changes occurring mainly on the protein level (Liguz-Lecznar et al., 2015). However, among the differentially expressed genes, we observed significantly diminished *Pvalb* transcript levels in 16 months old cortices (**Fig. 1b**), the time point when aging begins in mice (Xu et al., 2007) This data was confirmed by *in situ* hybridization experiments, indicating an age-related reduction of *Pvalb*-expressing cells in motor, somatosensory and visual neocortical areas (**Fig. 1c, d**). Consistently, we found less PV-immunoreactive cells (**Fig. 1e, f**) in the same cortical regions in aged mice. Together, our data suggest a loss of PV-positive cortical interneurons in aged mice, confirming the decrease of PV interneurons in somatosensory, auditory and motor cortical areas of aged rats (Miettinen et al., 1993; Ouda et al., 2008).

**Figure 1.**
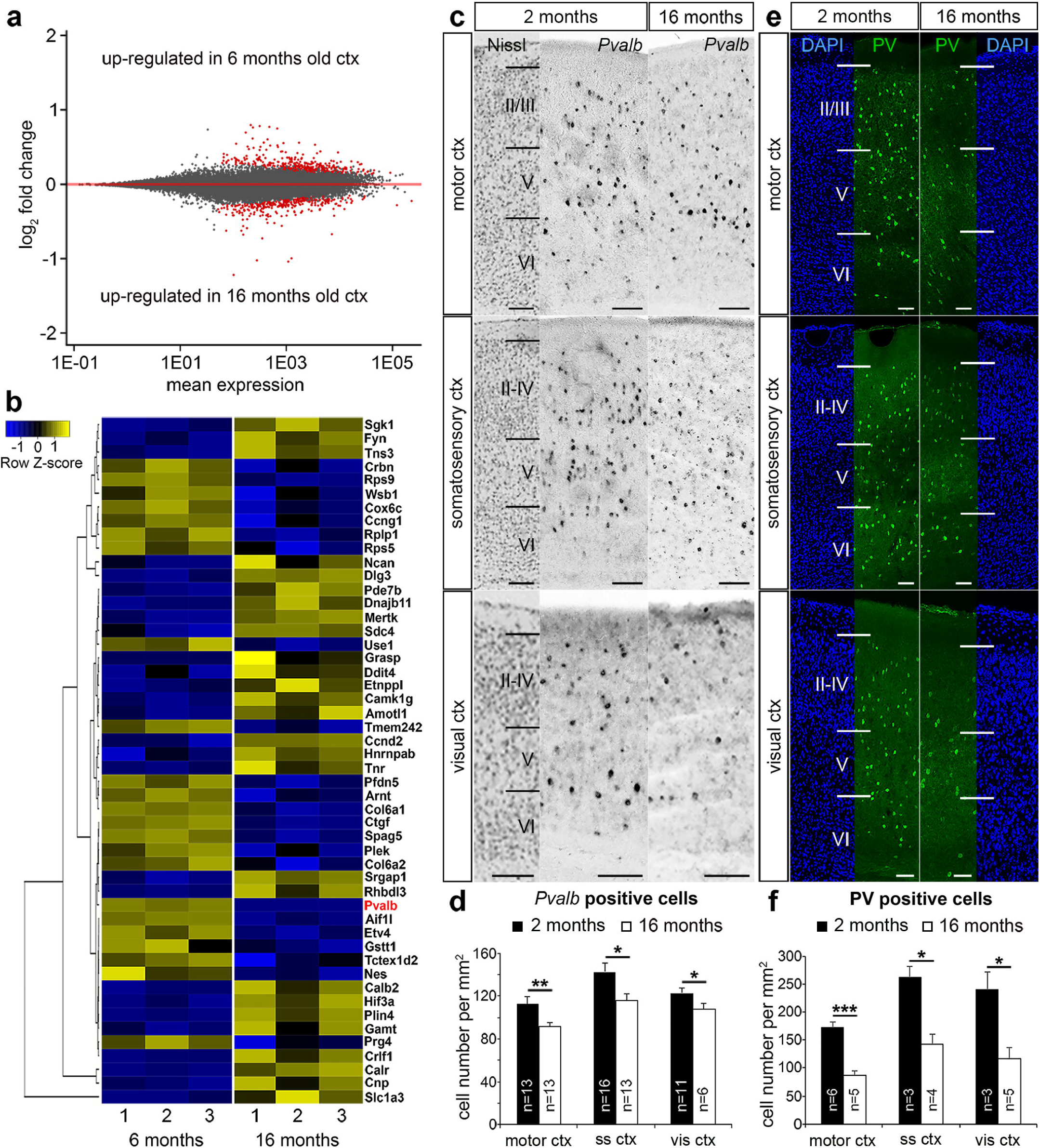
Age-dependent reduction of parvalbumin expression in the mouse cortex. (**a**) MA-plot illustrating differential gene expression between whole cortical tissue of 6 months (n = 3 brains) and 16 months old (n = 3 brains) C57BL/6 mice. RNA sequencing was performed to reveal age-dependent differential gene expression (red dots, *P* < 0.05, Benjamini adjusted), with negative ratios representing an up-regulation of gene expression in old cortical tissue samples. (**b**) Heat-map illustrating the 50 genes with highest fold-changes among the differentially expressed genes between cortical tissue samples of 6 and 16 months old C57BL/6 WT mice including *Pvalb*, which is significantly reduced in aged samples. (c-f) *In situ* hybridization and immunohistochemistry against *Pvalb* mRNA (c) and PV protein (e) in the motor cortex, somatosensory cortex and visual cortex in 6 months and 16 months old C57BL/6 mice (N = 3 different brains per age), quantified in (d) and (f), respectively (\**P* < 0.05; \*\**P* < 0.01; \*\*\**P* < 0.001, *Student’s t-test*). Scale bars: 100 µm in c and e

In contrast to this depletion of inhibitory PV interneurons in the cerebral cortex, excitatory neurons, which account for >80% of cortical neurons (DeFelipe & Fariñas, 1992), appear less affected. Pan-neuronal density analysis of NeuN-positive cells did not reveal significant changes upon aging (**Supplementary Figure S1**). In summary, we identified a vulnerability of PV-positive cortical inhibitory interneurons upon aging in mice.

### DNMT1 affects the long-term survival of neocortical interneurons

Changes of the epigenetic landscape by genomic methylation and histone modifications contribute to transcriptional control in aging and lifespan regulation (Zampieri et al., 2015). DNA methylation, executed by DNA methyltransferases (DNMTs), is a major epigenetic mechanism regulating gene expression in mammals during different stages of life (Johnson et al., 2012; Zampieri et al., 2015). DNMT1 is one of the main DNMTs expressed in the developing and adult brain. DNMT1 modulates neuronal survival (Feng et al., 2010; Hutnick et al., 2009; Pensold et al., 2017; Symmank & Zimmer, 2017) and synaptic function of both excitatory neurons (Meadows et al., 2015, 2016) as well as inhibitory interneurons (Pensold et al., n.d.). Hence, we asked whether DNMT1 is involved in the regulation of cortical interneuron survival during aging. To this end, we exploited a mouse model described previously (Pensold et al., n.d.), in which *Dnmt1* deletion is restricted to PV-cells (*Pvalb*-*Cre*/*tdTomato*/*Dnmt1 loxP*^*2*^). As controls, we used *Pvalb*-*Cre*/*tdTomato* mice. *Pvalb* promoter-dependent CRE recombinase-mediated *loxP* recombination drives tdTomato protein expression, reported to start at the 5^th^ week of life (Madisen et al., 2010). The analysis of the *Pvalb-Cre/tdTomato* interneuron density in adult *versus* aged mice confirmed the findings we obtained by RNA sequencing of whole cortical tissue, in situ hybridization, and immunostainings in C57BL/6 wildtype mice (**Fig. 1**). We found a significant age-related reduction of tdTomato positive cells in motor and visual cortical areas of *Pvalb-Cre/tdTomato* mice (**Fig. 2a, c**). Both superficial and deep cortical layers were affected by the reduction in interneurons (**Fig. 2d**). Although less prominent, we also observed a significant decline of tdTomato cells in the somatosensory cortex (**Fig. 2a, c**). This reduction was mainly restricted to the deep cortical layers (**Fig. 2d**). At an intermediate stage (12 months old mice) we found a trend for reduced cell density, indicating that interneuron degeneration starts about one year of life (**Fig 2a, c**).

**Figure 2:**
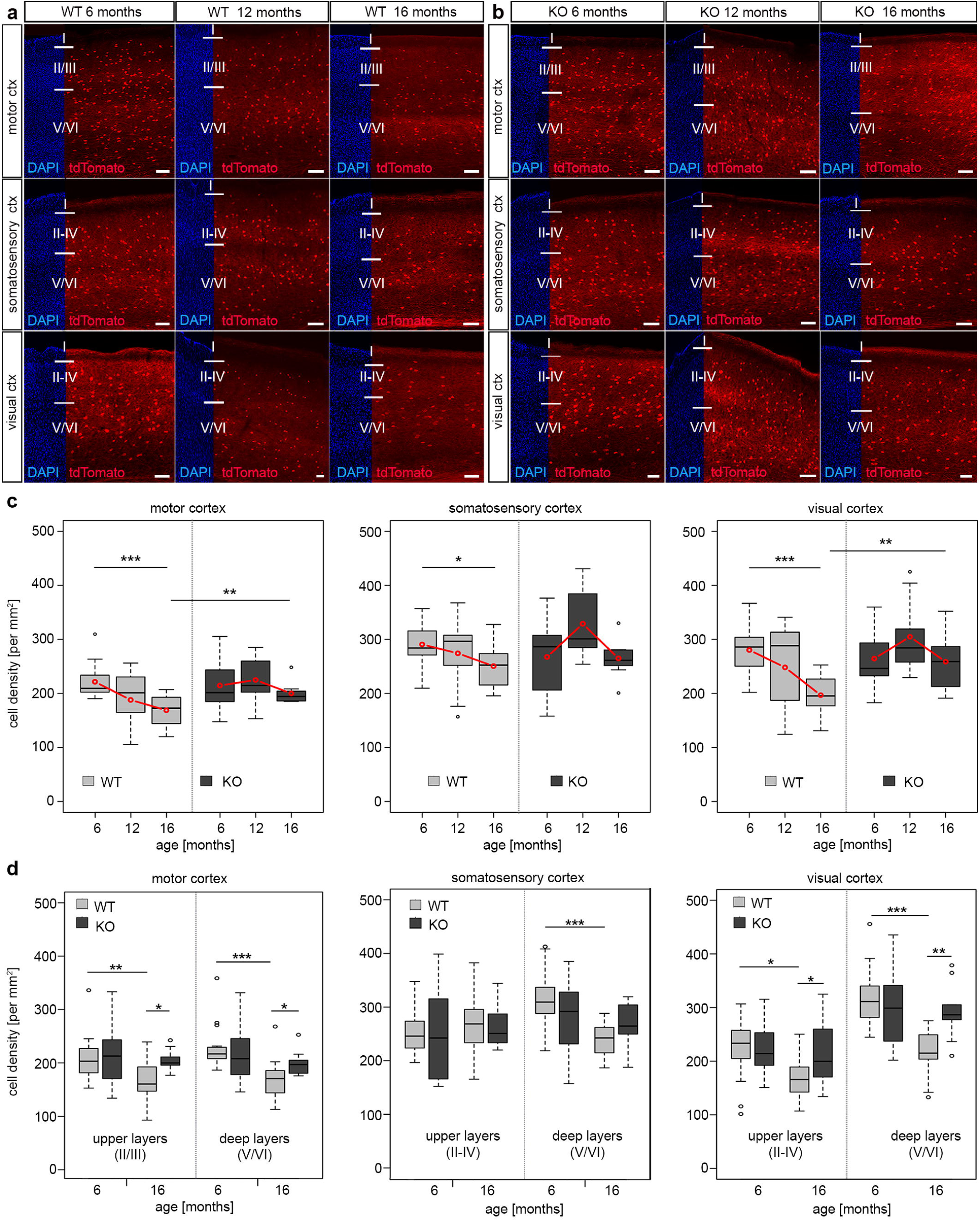
DNMT1 knockout enhanced long-term survival of cortical interneurons. (**a, b**) Microphotographs of sagittal sections (Bregma 1.44) illustrating the motor, somatosensory, and visual cortex of 6, 12 and 16 months old *Pvalb-Cre*/*tdTomato* (WT, **a**) and *Pvalb/tdTomato/Dnmt1 loxP*^*2*^ (KO, **b**) mice showing tdTomato (red) and DAPI positive cells (blue). The cell density per area is quantified in (**c**), which revealed a significant loss of interneurons upon aging in *Dnmt1* WT (\**P* < 0.05, \*\*\**P* < 0.001) for the motor, somatosensory and visual cortex, respectively (two-way ANOVA, Bonferroni corrected), but no significant age-dependent differences in *Dnmt1* KO mice. Comparison of aged genotypes revealed significant differences in the motor and visual cortex (\*\**P* < 0.01; *Student’s t-test*) (**d**) Layer-specific analysis of cell density in 6 and 16 months old *Dnmt1* WT and KO mice in the motor, somatosensory, and visual cortex (\**P* < 0.05, \*\**P* < 0.01; \*\*\**P* < 0.001, *Student’s t-test*). The numbers of analyzed sections are listed as follows: 6 months old WT: n = 16, n = 15 and n = 13 for motor, somatosensory and visual cortex, respectively; 6 months old KO: n = 19 for motor cortex and n = 16 for somatosensory and visual cortex; 12 months old WT and KO: n = 9 sections for each cortical area; 16 months old WT: n = 14 for motor and visual cortex and n = 12 for somatosensory cortex; 16 months old KO: n = 9 for each cortical region (from N = 3 different brains per genotype and age). WT = wild-type, KO = knockout. Scale bars: 100 µm.

Next, we comparatively analyzed tdTomato cells in 6 months, 12 months and 16 months old *Pvalb*-*Cre*/*tdTomato/Dnmt1 loxP*^*2*^ mice in the motor, somatosensory and visual cortical areas. While in young mice no differences in interneuron numbers were observed compared to controls (**Fig. 2b, c**), 16 months old *Dnmt1* KO mice maintained a significantly higher density of tdTomato positive interneurons in motor and visual cortical areas (**Fig. 2a-c**). In the somatosensory cortex, we again observed a trend towards increased densities of *Dnmt1*-deficient interneurons compared to age-matched controls **(Fig. 2a, c**). Hence *Dnmt1* deficiency substantially improves long-term survival of PV-expressing cortical interneurons, indicating that DNMT1 function either directly or indirectly impairs cortical PV-interneuron survival in aged mice. This is in striking contrast to DNMT1 function during brain development, where it promotes POA-derived interneuron survival through non-canonical actions (Pensold et al., n.d., 2017; Symmank et al., 2018).

### The ameliorated interneuron survival in aged *Dnmt1*-deficient mice correlates with improved somatomotor performances

Given their important role in cortical information processing, cortical interneuron decline was proposed to contribute to the cognitive decline and motoric impairments observed in the elderly (Bordner et al., 2011). To test whether attenuated interneuron loss correlates with improved skills in aged *Dnmt1*-deficient mice, we applied the ladder rung test to analyze motor performance that depends on somatomotor cortical activity (Metz & Whishaw, 2009). We continuously tested *Pvalb*-*Cre*/*tdTomato*/*Dnmt1 loxP*^*2*^ and *Pvalb*-*Cre*/*tdTomato* mice at distinct stages of life ranging from 3 to 21 months. Consistent with observations of others (Hebert & Gerhardt, 1998) and the age-dependent changes in interneuron numbers, the motor performances of control mice deteriorated with age as determined by measuring the foot placement accuracy and crossing time (**Fig. 3a-c**). In stark contrast, *Dnmt1*-deficient mice did not show corresponding age-related impairments for the parameters and the time course analyzed, hence performing significantly better than controls at 16 to 21 months of age (**Fig. 3a-c**). When plotting the percentage of perfect steps against crossing time for KO and control mice at 6, 12, and 18 months (**Fig. 3d-f**), cohort segregation increased with age.

**Figure 3.**
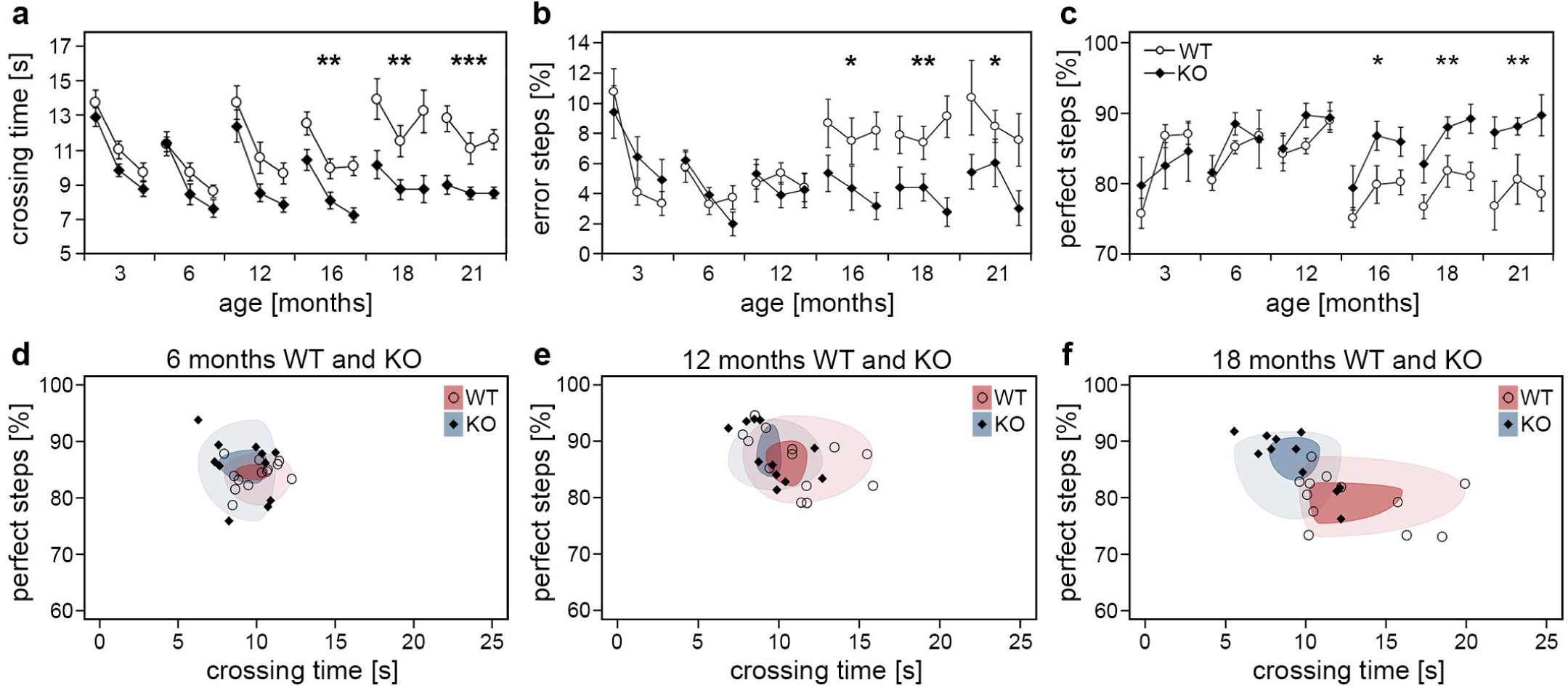
Aged *Pvalb*-Cre/*tdTomato*/*Dnmt1* KO mice show improved somatomotor performances. Performance of *Pvalb*-Cre/*tdTomato*/*Dnmt1* KO (N=10) and WT (N=8) mice were tested in the inclined ladder rung test at distinct stages of life ranging from 3 to 21 months (three consecutive trials per stage). Crossing time (**a**), error steps (**b**) and perfect steps (**c**) were quantified (two-way ANOVA; data are shown as mean ± SEM; \**P* < 0.05, \*\**P* < 0.01, \*\*\**P* < 0.001). (**d**-**f**) The scatter plots of perfect steps against crossing time for the distinct *Pvalb*-Cre/*tdTomato*/*Dnmt1* KO and WT mice at 6 months (**d**), 12 months (**e**) and 18 months (**f**) illustrate the stronger segregation of the cohorts with age shown by the decreasing overlap of the circles upon aging representing the 1^st^ (dark colored) and 3^rd^ quartile (light colored) of the data range per group.

In addition to cortical information processing, locomotion depends on cerebellar Purkinje cells and skeletal muscle function, tissues that also display *Pvalb* and *Dnmt1* expression (**Fig. S2a, b;** Racay et al., 2006). In skeletal muscle, DNMT1 indeed plays a role during differentiation and regeneration (Aguirre-Arteta et al., 2000; Wang et al., 2015). However, neither in skeletal muscle nor in the cerebellum, obvious abnormalities were observed upon *Dnmt1* deletion. Purkinje cell numbers in the cerebellum were not affected by PV-CRE mediated *Dnmt1* deletion, neither in the young nor in the aged mice (**Fig. S2c-e**). Moreover, muscle integrity, structure, and innervation were not altered by *Dnmt1*-deletion at the stages investigated, as determined by hematoxylin/eosine, laminin and neuromuscular junction staining, respectively (**Fig. S2f-k**). These data strongly suggest that the motor impairments in aged controls are caused by the loss of cortical interneurons, which can be attenuated by *Dnmt1* deletion.

### PV interneurons show an increase in degradation- and a decline in synapse-related gene expression upon aging

Highlighting age-mediated transcriptional changes might help to approach the underlying mechanisms of the DNMT1-dependent PV-interneuron loss. This requires enrichment of PV-positive cortical interneurons from adult *versus* aged brains, as these interneurons represent a minority of the neocortical neuronal population (Druga, 2009). To this end, we applied an optimized protocol for adult cortical neuron isolation applicable for fluorescence activated cell sorting (FACS). We combined mechanical and collagenase-based enzymatic dissociation with trehalose treatment and *Percoll* density gradient centrifugation, as described and validated recently (Pensold et al., n.d.).

Previously, Xu et al. (2007) investigated murine brain tissue at 6, 16 and 24 months of age, and found that most age-dependent genes are not differentially expressed at the age of 16 months. Hence, we chose to analyze interneurons of 18 months old control *versus* conditional *Dnmt1* knockout mice to monitor an advanced stage of aging, and compare interneuron transcriptional profiles with 6 months old mice for each genotype. Consistent with the PV interneuron loss in aged controls, we revealed significantly reduced FACS-events per hemisphere for aged *Pvalb*-*Cre*/*tdTomato* mice compared to the 6 months old mice (**Supplementary Figure 3a**). Transcriptome comparison between FAC-sorted young and old control interneurons illustrated that aging is associated with prominent changes in gene expression (**Fig. 4a, Supplementary Figure 3b-d**). 3384 genes were differentially regulated (adjusted p<0.05, | log2fc |>1), of which 65% were down-regulated and 35% up-regulated with age (**Fig. 4a**). This high number of age-dependent transcriptional changes exceeds the transcriptional alterations revealed for the whole cortex (**Fig. 1a, b**), which captures different aging signatures of diverse cell types collected in the cortical samples (Kimmel et al., 2019; Stegeman & Weake, 2017). Among the genes we found to be up-regulated upon aging, gene ontology (GO)-enrichment analysis revealed significantly enriched transcripts related to *membrane, endoplasmatic reticulum, endosome* and *exosome* (**Supplementary Table S1**). The up-regulation of endosome and exosome-related genes in cortical interneurons might reflect an elevation of degradative actions and mechanisms upon aging in response to the accumulation of defective proteins, to maintain neuronal functionality over life time.

**Figure 4.**
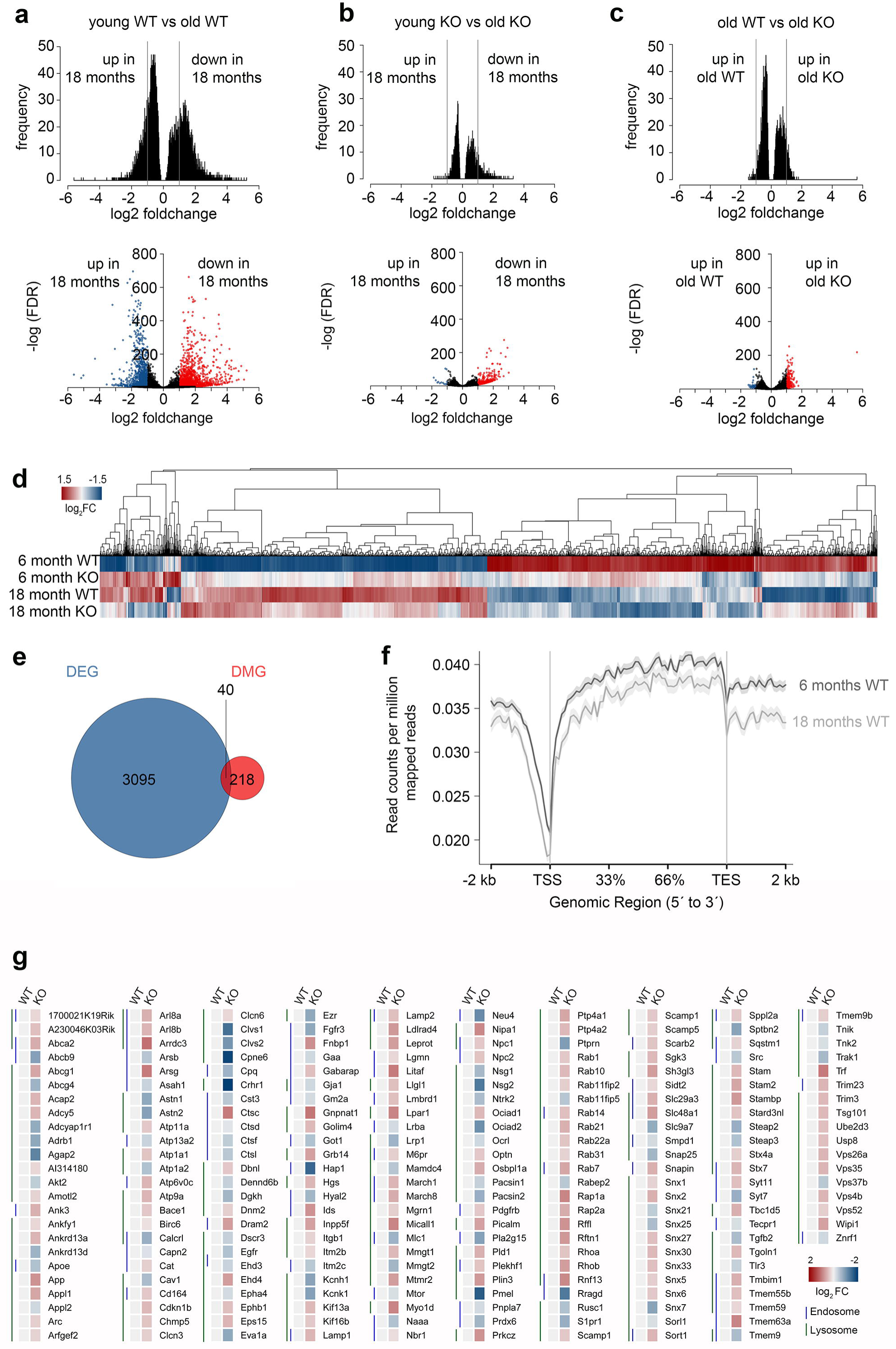
Correlative transcriptome and methylation analysis of adult and aged *Dnmt1*-deficient and wild-type *Pvalb*-expressing cortical interneurons. (**a**-**c**) Density plots (upper panel) and volcano plots (lower panel) illustrating significant changes in gene expression determined between FACS-enriched young and old *Dnmt1* WT interneurons (**a**), young and old *Dnmt1* knockout (KO) cortical interneurons (**b**), as well as between old WT and KO interneuron samples (**c**; *P* < 0.05, Benjamini adjusted; pooled samples from N = 6 WT and KO mice at 6 months; and N = 9 WT and N = 12 KO mice at 18 months analyzed in technical duplicates). Blue and red-colored dots in the volcano plots represent genes with fold change > 2. (**d**) Heat-map illustrating the re-scaled expression of genes in all samples, which were found differentially expressed between young (6 months) KO and young WT interneurons. (**e**) Venn diagram illustrating the non-significant overlap (*P*=3.388E-5, *Fisher’s Exact test*) of differentially expressed (DEG) and differentially methylated genes (DMG) between aged FACS-enriched *Pvalb*-Cre/*tdTomato*/*Dnmt1* WT and KO cortical interneurons as determined by RNA-sequencing (*P* < 0.05, Benjamini adjusted) and MeDIP-sequencing (n = 9 WT and n = 12 KO mice; *P* < 0.05, Benjamini adjusted). (**f**) Methylation plot illustrating the average DNA methylation levels of a random sample of 10% of the genes from the *mm10* reference genome in young (6 months) and old (18 months) cortical interneurons from *Dnmt1* WT mice. (**g**) Heat-map of differentially expressed genes associated to the GO terms *endosome, autophagy* and *lysosome*, normalized to 6 months old WT.

Of note, functional impairment of exosomes in transferring proteins, mRNAs and miRNAs has been related to synaptopathies (Pitt et al., 2017), and synaptic dysfunction is considered a hallmark in neuronal aging (Azpurua & Eaton, 2015; Deak & Sonntag, 2012) and neurodegenerative disorders (Freeman & Mallucci, 2016; Ghiglieri et al., 2018). Altered or impaired synaptic function of aged PV-expressing interneurons is strongly supported by the profile of genes that were down-regulated upon aging. By GO analysis, synapse-related genes were detected as most significantly overrepresented, displaying by far the highest enrichment scores (Benjamini= 1.91E-61; FDR=4.6E-61; Table S1). Moreover, genes collected in the GO-terms *membrane, cell junction, plasma membrane, dendrite* and diverse ion transport and ion channel-related genes were strongly enriched among the genes determined as transcriptionally down-regulated in aged wild-type interneurons (**Supplementary Table S1**). In sum, the transcriptional alterations that we detected in aged neocortical PV-positive interneurons suggest an age-related impairment of synaptic functionality. Moreover, alterations in the degradation machinery can be assumed from the transcriptional alterations, which can influence neuronal survival (Kim & Seo, 2014).

### *Dnmt1* deficient PV interneurons display diminished age-associated transcriptional alterations

In addition to ameliorated locomotion, the attenuated decline in interneuron density in aged *Dnmt1* knockout mice coincides with diminished age-associated quantitative transcriptional changes in *Pvalb*-*Cre*/*tdTomato*/*Dnmt1 loxP*^*2*^ interneurons (**Fig. 4b; Supplementary Table 2**). Compared to control interneurons, aging in *Pvalb*-*Cre*/*tdTomato*/*Dnmt1 loxP*^*2*^ cortical interneurons was characterized by both fewer differentially expressed genes and decreased fold changes. Only 383 genes were differentially expressed (adjusted p<0.05, | log2fc |>1; F**ig. 4a, b**). For better illustration of the discrete changes in expression between all samples, we re-scaled the expression levels of genes relative to the expression range of all groups (young and old control as well as knockout samples; **Fig. 4d**). The heatmap shown in **Fig. 4d** depicts prominent age-related transcriptional alterations in controls, but rather mild alterations in *Dnmt1*-deficient interneurons. These data are consistent with the attenuated age-associated decline observed for conditional *Dnmt1*-knockout mice at cellular and behavioral level.

A common denominator of age-mediated transcriptional remodeling in both genotypes is that age-related down-regulation dominates over up-regulation for the significantly altered genes with | log2fc |>1 (**Fig. 4a, b**). For age-associated gene expression changes in *Pvalb*-*Cre*/*tdTomato*/*Dnmt1 loxP*^*2*^ interneurons, about 96% of differentially expressed genes were down-regulated (**Fig. 4b**). Consistently, a “shutdown” of transcription in the aged cortex has been described before (Xu et al., 2007). Another similarity between aging control and *Dnmt1*-knockout interneurons was a significant enrichment of down-regulated synapse-related genes (**Supplementary Table S1, S2**).

### DNA methylation plays a minor role in age-mediated transcriptional remodeling

DNA methylation was frequently proposed to contribute to the aging-dependent transcriptional changes (Issa, 2002; Jones et al., 2015). To this end, we conducted differential methylation analysis by MeDIP-sequencing of FAC-sorted interneurons from young (6 months) and aged (18 months) control mice. Next, we determined genes whose age-related transcriptional changes (adjusted p<0.05) correlated which alterations in the DNA methylation level (adjusted p<0.05). Among the 201 genes which demonstrated changes in expression and DNA methylation upon aging, *synapse, cytoskeleton, dendrite, postsynaptic density* and *membrane*-related genes were significantly overrepresented (**Supplementary Table 3**), indicating that DNA methylation contributes to the age-related transcriptional changes of these genes.

To determine which genes are differentially expressed and methylated in aged interneurons in a DNMT1-dependent manner, we compared transcriptional profiles and DNA methylation signatures of old control and *Dnmt1*-deficient interneuron samples. We obtained only 258 differentially expressed genes (adjusted p<0.05) displaying a | log2fc |>1 **(Fig. 4c)**. 218 genes showed differential methylation (adjusted p<0.05), of which only 2 genes were overlapping with the differentially expressed genes. Hence, we included all significantly differentially expressed genes independent of their fold change (3095 genes) for correlation analysis between changes in methylation and transcription. Only 40 genes were significantly changed in both expression and methylation between the aged genotypes (**Fig. 4e**). This represents a non-significant overlap (*P*=3.388E-5, *Fisher’s Exact test*), indicating that DNA methylation rather plays a minor role for in transcriptional control of aged brains. Indeed, the efficiency of the catalytic activity of DNMT1 is described to be reduced in an age-dependent manner (Casillas et al., 2003). This is in line with the global reduction of DNA methylation levels observed upon aging in control interneurons (**Fig. 4f**), a finding that corroborates the age-related global hypomethylation reported by others (Lardenoije et al., 2015; Shimoda et al., 2014).

For those 40 genes which simultaneously changed in both expression and DNA methylation between aged genotypes, GO analysis revealed a significant enrichment of *actin cytoskeleton* and *postsynaptic density*-related genes, which are putatively regulated by DNMT1-dependent DNA methylation in aged interneurons (**Supplementary Table 4**). This fits to our finding that synapse and cytoskeleton-related genes are DNA methylation-dependently changed in expression upon aging (**Supplementary Table 3**). However, DNA methylation by DNMT1 seems to have only minor a impact on transcriptional regulation in aged interneurons. Therefore, DNMT1 target genes identified in 6 months old mice hold more promise to identify the causes of impaired long-term survival.

### DNMT1-dependent DNA methylation in young adult interneurons affects degradative pathways

In stark contrast to aged genotypes, we determined a significant overlap (*P* = 2.2E-16, *Fisher’s exact test* for gene set enrichment analysis; odds ratio = 0.434) of 645 genes between young control and *Dnmt1* knockout interneurons. These 645 genes display significant differences in both DNA methylation and gene expression (Pensold et al., n.d.).

In general, far more genes were differentially expressed (3868 genes) and / or methylated (3135 genes) between young genotypes (Pensold et al., n.d.). However, among neither the differentially expressed genes, nor among those genes both differentially expressed and methylated, we found a significant enrichment of apoptosis or cell death-related genes (data not shown). Hence, in contrast to developing interneurons, where DNMT1 regulates expression of apoptosis genes (Pensold et al., 2017), survival regulation of interneurons in the aged cortex seems to result from different actions and targets of DNMT1.

Among the genes which we identified as repressed by DNMT1-dependent DNA methylation in young controls, we found an overrepresentation of *endocytosis* and *endosome*-related genes (Pensold et al., n.d.). Furthermore, during analysis of all genes differentially expressed upon *Dnmt1* deletion, irrespective of altered DNA methylation, *lysosome* and *ubiquitination*-related genes were also found repressed by DNMT1 (Pensold et al., 2020; **Fig. 4g**). Together these results demonstrate that endocytosis and degradative pathways are regulated by DNMT1. In a previous study we confirmed that dynamic DNMT1-dependent DNA methylation regulates synaptic transmission through the modulation of endocytosis-mediated vesicle recycling, which was improved upon *Dnmt1* deletion (Pensold et al., n.d.). Hence, elevated GABAergic transmission and synaptic activity could indirectly promote interneuron survival of *Dnmt1*-deficient interneurons upon aging. However, endocytosis and endosomal function are crucial not only for synaptic activity regulation, but also affect degradative pathways (Ehlers, 2000; Gruenberg, 2001). Consistent with the transcriptional changes in 6 months old *Dnmt1*-deficient cortical interneurons (**Fig. 4g**), siRNA-mediated *Dnmt1* depletion (knockdown efficiency of *Dnmt1* siRNA is illustrated in **Supplementary Fig. 4a**) caused augmented CD63 and LAMP1 immunoreactivity, labeling endosomal and lysosomal structures, respectively. This was determined in neurites of interneurons prepared from the embryonic medial ganglionic eminence (**Fig. 5a-c**) that give rise to PV interneurons, as well as in neurite-like processes of cerebellar granule (CB) cells and neuroblastoma N2a cells 24 hours after transfection (**Supplementary Fig. 4b-d**).

**Figure 5:**
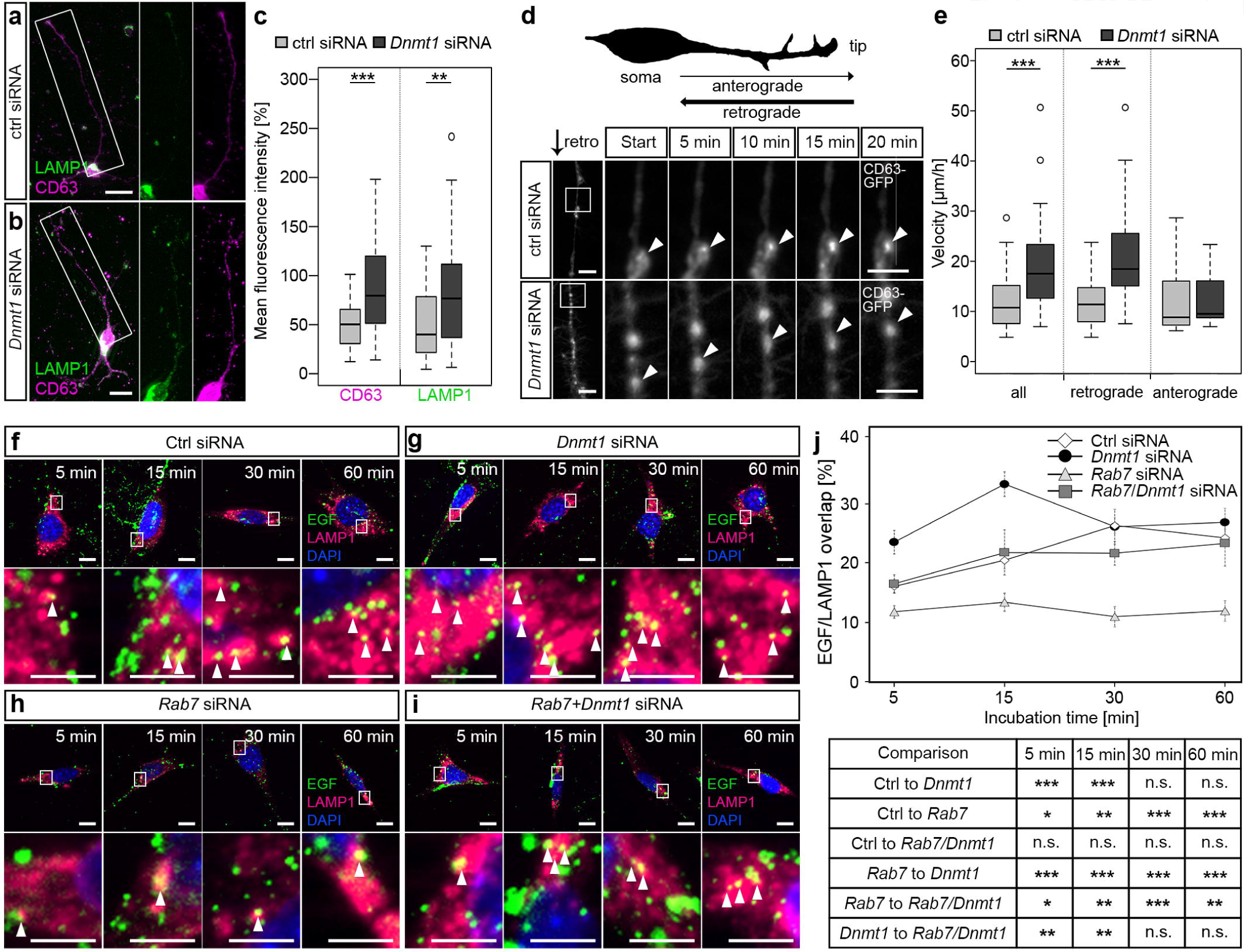
DNMT1 regulates retrograde transport of endosomes and endocytic-based degradation. (**a, b**) CD63 (magenta) and LAMP1 (green) antibody staining in MGE cells (E15+7 div) 24 h after control and *Dnmt1* siRNA treatment. The white rectangles in (**a, b**) illustrate the locations of the magnified parts of the processes. Quantification of fluorescence intensities is shown in (**c**). (**d, e**) Cerebellar granule (CB) cells were co-transfected either with control or *Dnmt1* siRNA and a CD63-GFP expression plasmid, and the movement of CD63-GFP positive structures was imaged for 20 min. The location of the highly magnified sections along the neurite-like processes are illustrated by white squares in the low magnifications. (**d**) Schematic illustration of the morphology of a cultured cerebellar granule (CB) cell. The quantification is depicted in (**e**). (**f-j**) The EGF-degradation assay with CB cells either transfected with (**f**) control siRNA, (**g**) *Dnmt1* siRNA, (**h**) *Rab7* siRNA or (**i**) *Dnmt1* and *Rab7* siRNA. The co-localization of Alexa488-labeled EGF (green) and LAMP1 positive lysosomal structures (red) was quantified after 5 min, 15 min, 30 min and 60 min, as quantified in **j**. CB cells = cerebellar granular cells; N2a cells = neuroblastoma cells. *Student’s t-test* in **c, e** and **j** with \**P* < 0.05, \*\**P* < 0.01, \*\*\**P* < 0.001. Scale bars: 20 µm in (**a, b**); 10 µm in (**d, f-i**); 5 µm in magnified sections in (**d, f-i**).

Endosomal-based degradation involves ubiquitination, retrograde transport to the cell soma, and fusion with lysosomes (Haglund & Dikic, 2012; McMahon & Boucrot, 2011). To quantify whether retrograde shuttling of endosomal compartments is influenced by DNMT1, we transfected CB cells with a CD63-GFP construct and analyzed transport velocity in the neurite-like processes upon *Dnmt1* knockdown and control siRNA transfections. While anterograde transportation was not changed in speed, we determined significantly faster velocities for retrograde transport of CD63-GFP particles after *Dnmt1* depletion (**Fig. 5d, e; Supplementary Movies 1-2**).

As the binding of epidermal growth factors (EGF) to epidermal growth factor receptors (EGFR) induces their internalization and degradation via the endo-lysosomal pathway (Haglund & Dikic, 2012), we next applied Alexa488-coupled EGF to CB cells and monitored the co-localization of EGF with LAMP1-positive lysosomal compartments at different time points. Indeed, siRNA-mediated *Dnmt1* depletion caused increased co-localization of EGF with LAMP1-positive lysosomes, both 5 and 15 min after EGF application (**Fig. 5f, g, j**), indicating transport to lysosomal compartments. With longer incubation times, no differences to control siRNA-treated cells were observed (**Fig. 5f, g, j**).

Lysosomal trafficking of the EGF-EGFR complex depends on RAB7, which mediates the fusion of late endosomes with lysosomes (Bucci et al., 2000). Consistently, we revealed a reduced EGF/LAMP1 co-localization after *Rab7* siRNA transfection of CB cells at all time points tested (**Fig. 5h, j**). *Rab7* expression was significantly up-regulated in *Dnmt1*-deficient PV-positive cortical interneurons (**Fig. 4g**), and shown to be regulated by DNMT1-dependent DNA methylation (Pensold et al., n.d.). Thus, we additionally analyzed the EGF/LAMP1 co-localization in *Dnmt1* siRNA-treated CB cells that were co-transfected with *Rab7* siRNA (knockdown efficiency of *Rab7* siRNA is depicted in **Supplementary Figure 4a**) to counteract the gain in *Rab7* expression in *Dnmt1*-siRNA transfected cells. This reversed the *Dnmt1* siRNA-triggered increase in EGF/LAMP1 co-localization (**Fig. 5i, j**), suggesting that DNMT1 restricts endocytic-based degradation partly through repression of *Rab7* expression.

Ubiquitination is a common denominator in the targeting of substrates to the main protein degradation pathways (Clague & Urbé, 2010), including lysosomal degradation (reviewed in Clague & Urbé, 2006). Interestingly, we determined elevated proportions of ubiquitin-positive cortical interneurons evident in *Dnmt1*-deficient mice (50 ± 0.8%) compared to wild-type controls (39.5 ± 2%; \*\**P* < 0.01, *Student*’s t-test; n = 3 mice per genotype; **Supplementary Fig. 4e, f**). Together, our data indicate that DNMT1 acts repressive on intracellular degradative pathways, which could affect long-term neuronal survival.

## 4 Discussion

We here provided evidence that DNMT1 is implicated in the compromised long-term survival of inhibitory PV interneurons in the murine cerebral cortex. Aging is characterized by reduced PV interneuron numbers accompanied by a decline in somatomotor performances and prominent transcriptional remodeling. All of these effects were attenuated by *Dnmt1* deletion in PV interneurons. While DNMT1 promotes neuronal survival in the developing nervous system, it seems to compromise the long-term survival of PV-interneurons in the aged cortex. Our global transcriptome and methylome analyses suggest that DNA methylation-dependent actions of DNMT1 in the aged interneurons seem to play, if any, only a subtle role in transcriptional regulation. As repressive DNMT1-dependent DNA methylation restricts synaptic transmission as well as degradative pathways in young PV interneurons, we hypothesize that the impaired long-term survival is an indirect consequence of DNMT1-mediated modulation of synaptic activity and degradation over life-time.

### Vulnerability of inhibitory cortical interneurons towards aging

Besides reduced excitability and plasticity (Clark & Taylor, 2011) and a decline of inhibitory function (Cheng & Lin, 2013; Shetty & Turner, 1998; Stanley & Shetty, 2004), selective vulnerability of particular neuronal subtypes, like inhibitory interneurons, and GABAergic synapses (Rozycka & Liguz-Lecznar, 2017) was reported for brain aging. Indeed, given the crucial role GABAergic inhibitory interneurons have in cortical information processing, age-dependent defects in inhibitory circuits provide an attractive hypothesis for the cognitive decline and age-associated disorders (Rozycka & Liguz-Lecznar, 2017).

In agreement with our finding that PV interneuron numbers were reduced in old cortices, several studies reported declined cell numbers of cortical interneuron subtypes across different species and brain regions (reviewed in Zimmer-Bensch, 2019). Several studies across mammalian species described an age-related decrease of glutamate decarboxylase-67 (GAD-67)-positive interneurons in the hippocampus (Shetty & Turner, 1998; Shi et al., 2004; Stanley & Shetty, 2004; Vela et al., 2003) and other cortical areas (Ling et al., 2005; Stranahan et al., 2012). Apart from the PV interneuron subpopulation, SOM, CB, VIP and NPY-positive cells appear to be affected by the aging process across diverse cortical areas (reviewed in Zimmer-Bensch, 2018). Moreover, functional and structural changes of GABAergic synapses appear to occur in aged brains, including the loss of synaptic contacts, a decreased neurotransmitter release, and a reduced postsynaptic responsiveness to neurotransmitters (Rozycka & Liguz-Lecznar, 2017). In agreement with reduced neurotransmitter release, major changes in the expression of genes related to neurotransmission and transcriptional repression especially of GABA-related transcripts have been reported for the human prefrontal cortex (Loerch et al., 2008). Consistently, age-related changes in mRNA or protein levels of GABA-associated genes were reported across different species (reviewed in Rozycka & Liguz-Lecznar, 2017; Zimmer-Bensch, 2019).

Surprisingly, we found that *Dnmt1* deletion significantly improves PV-interneuron survival in aged motor and visual cortices of mice, indicating that DNMT1 is implicated in the age-related interneuron loss.

### DNMT1-dependent survival regulation in the nervous system

Despite reports of DNMT-dependent developmental regulation of neuronal survival (Hutnick et al., 2009; Pensold et al., 2017; Rhee et al., 2012), evidence for direct survival regulation by DNMTs in the aging brain is still lacking. Support for potential functional implications of DNA methylation in neuronal cell death regulation emerged from patients diagnosed with Alzheimer’s Disease (AD), an aging-associated neurodegenerative disorder. In neurons from postmortem cortical tissue 5mC and 5hmC immunoreactivity was found to be globally altered compared to age-matched control individuals (Coppieters et al., 2014; Mastroeni et al., 2010).

In case DNMT1 would act on transcriptional networks implicated in neuronal survival regulation in PV interneurons, differences in expression profiles of survival-, apoptosis- or cell death-related genes should be evident between adult and/or aged *Dnmt1*-deficient and control interneurons. However, we found no enrichment in apoptosis, cell death or cell survival regulating genes, neither in differential gene expression analyses between young nor between aged genotypes. The same is true for expression profiles of young and aged control interneurons. Hence, the comparative gene expression analysis among PV interneuron populations strongly implies that in contrast to developing interneurons, DNMT1 does not affect their long-term survival in the aging brain by the transcriptional control of survival and/or cell death-related genes.

### Potential implications of DNMT1-mediated regulation of synaptic function

A decline in synaptic density and functionality, reduced synaptic markers, pruning of the dendritic tree, loss of dendritic spines, structural changes within the presynaptic active zone and alteration of receptors for different neurotransmitters have been detected across the nervous system in physiological aging (Berchtold et al., 2013; Burke & Barnes, 2006; Polydoro et al., 2009; Tanaka et al., 1996). Reduced densities were also seen for inhibitory synapses in the cerebral cortex (Calì et al., 2018). In line with the findings of synaptic impairments in the aging brain, we found that synapse-related genes were downregulated in old PV interneurons (**Table S1**). Such a decline in synaptic gene expression upon aging was already reported by others (Berchtold et al., 2013; Dillman et al., 2017; Loerch et al., 2008). In controls, the age-mediated transcriptional changes of synapse-associated genes correlate with alterations of the DNA methylation level (**Table S3**). Some of these synapse-related genes appear to be regulated by DNMT1-dependent DNA methylation, as concluded from the comparison of old control and old *Dnmt1* knockout samples (**Table S4**). This proposes an age- and DNMT1-dependent shutdown of synapse-associated gene expression, which could impair synaptic function. As activity-dependent signaling is described to boost neuronal health through diverse mechanisms, decreased synaptic functionality could indirectly act on neuronal survival. Besides transcriptional control of pro- and anti-apoptotic genes, the availability of neurotrophic factors and the elevation of antioxidant defenses is modulated by neuronal activity (reviewed in Bell & Hardingham, 2011). In this context, it is not surprising that activity-dependent signaling influences multiple aspects and events of individual neurodegenerative diseases (reviewed in Bell & Hardingham, 2011).

We have recently shown that DNMT1 acts on synaptic function of cortical PV interneurons in young mice, modulating GABAergic transmission (Pensold et al., n.d.). Alterations in transmitter release affect synaptic strength, both of which is found to be decreased upon aging (Kumar et al., 2007). Hence, it could be conceivable that the increased rates of synaptic transmission seen in young *Dnmt1*-deficient interneurons have a protective effect on age-associated synaptic impairments, and thereby indirectly promote the survival in aged *Dnmt1*-deficient mice.

### DNMT1-dependent regulation of proteostasis and implications for neuronal aging

The long-term health of neurons ultimately depends on the interplay of the proteostasis network components. An age-related decline in protein homeostasis can cause diverse cellular dysfunctions and neuronal death, thereby contributing to numerous neurodegenerative disorders (Douglas & Dillin, 2010). An important finding made in this study is that genes related to the proteostasis network, like *membrane*-, *exosome*-, *endoplasmatic reticulum*-, and *endosome*-associated genes were upregulated upon aging.

Endosomal-based degradative pathways are crucial for processing and removing defective proteins or protein aggregates, either through proteolytic degradation in lysosomes (McMahon & Boucrot, 2011), or by exosomal-based release (Riva et al., 2019). Exosomal release is fundamental for protein removal into the extracellular space by inclusion into multivesicular bodies (MVBs) (Riva et al., 2019). Albeit the role of exosomes in normal neuronal aging is still under debate, exosomes have been recently implicated in contributing to neurodegenerative disease and mental disorders (Howitt & Hill, 2016; Kalani et al., 2014; Schneider & Simons, 2013).

Lysosomes are implicated in the digestion of extracellular material taken-up by endocytosis, as well as of intracellular material segregated by autophagy (Stoka et al., 2016). Lysosomal degradation appears compromised in aged neurons, while this is still discussed controversially in the literature (reviewed in Loeffler, 2019). Implications of lysosome-dependent lifespan regulation rely on their fundamental role in autophagy. The connection between autophagy and longevity is based on diverse observations. For example, suppression or loss of autophagy in the central nervous system causes neurodegenerative disease in mice (Hara et al., 2006; Komatsu et al., 2006). Moreover, maintaining high levels of autophagy is suggested to contribute to longevity (Triplett et al., 2015). Mice lacking *Atg7* (autophagy related 7), encoding for the E1-like activating enzyme that is essential for autophagy (Komatsu et al., 2005), develop neuronal loss and die within 28 weeks (Komatsu et al., 2006). In *C. elegans*, the knockdown of *Atg7* and *Atg12* shortens the lifespan (Hars et al., 2007), illustrating the relevance of the proteostasis network for neuronal survival. Besides executing proteolytic degradation of cargo delivered by autophagic or endocytotic pathways, lysosomal function in turn executes regulatory control over autophagy. Lysosomes control the docking of autophagy-regulating factors like mTORC1 or TFEB to the lysosomal membrane, which determines their functionality (reviewed in Settembre et al., 2013).

Apart from their role for disposal and processing of cellular waste, lysosomes act as pivotal regulators of cell homeostasis at multiple levels, being suggested as a central cellular hub for aging control. Lysosomes are implicated in the regulation of cellular responses to nutrient availability and composition, stress resistance, programmed cell death, and plasma membrane repair (Boya, 2012; Braun et al., 2015a; Braun et al., 2015b; Settembre et al., 2013). Given this pleiotropic importance of lysosomes, it is not surprising that their dysfunction is associated with aging as well as with a plethora of neurodegenerative disorders (Jiang et al., 2001), and age-related pathologies like Parkinson’s and Alzheimer’s disease (Büttner et al., 2013; McBrayer & Nixon, 2013; Menezes et al., 2015; Wolfe et al., 2013).

As a declining proteostasis network accompanies aging and triggers ineffective protein degradation, the aggregation of defective proteins finally can lead to cell death (Douglas & Dillin, 2010). Hence, the up-regulation of genes related to proteostasis in control interneurons might be interpreted as a compensating response of aging neurons to counteract the remittent proteostasis network (Douglas & Dillin, 2010). This is in line with findings of Rodríguez-Muela et al. (2013), reporting an age-related increase in LAMP-2a and HSPA8/Hsc70 concentrations in mouse retina, suggested to compensate for the age-related decrease in macroautophagy. An age-dependent elevation of HSPA8/hsc70 levels was also seen in the hippocampus, cortex, cerebellum, septum and striatum (Calabrese et al., 2004).

Interestingly, such increase in proteostasis-associated gene expression was not seen upon aging in *Dnmt1*-deficient interneurons. This can be explained by the finding that *Dnmt1* deletion itself acts on proteostasis-associated gene expression in young interneurons. Compared to equal-aged controls, *endocytosis-, endosome-* and *lysosome*-related genes were increased in expression in *Dnmt1*-deficient samples (Pensold et al., 2020, **Fig. 4g**). While we previously verified that endocytosis-mediated elevated vesicle recycling increases the GABAergic transmission of *Dnmt1*-deficient interneurons, DNMT1-dependent regulation of degradative pathways remained unattended so far. In the present study, we validated that *Dnmt1* depletion elevated retrograde endosomal transport and lysosomal targeting, suggesting an improved degradative machinery upon *Dnmt1*-depletion. Such boosted degradative actions could be neuroprotective or beneficial for neuronal survival in the long run, preventing age-related interneuron loss as seen in *Dnmt1*-deficient mice.

Together, our data suggest that the DNA methylating activity of DNMT1 appears to play a rather minor role in age-related transcriptional remodeling. We identified only few genes displaying differential methylation and expression between the aged genotypes, as well as globally reduced DNA methylation levels upon aging in wild-types (**Fig. 4e, f**). This is consistent with the aging-associated global hypomethylation (Lardenoije et al., 2015; Shimoda et al., 2014) and age-dependent reduction in DNMT1 efficiency (Casillas et al., 2003). We rather anticipate that DNMT1-dependent changes in aged interneurons are a cumulative effect of its function during life-time, as DNMT1 modulates two crucial aspects for neuronal function: synaptic activity and proteostasis. Hence, we propose a scenario, in which *Dnmt1* deficiency-induced enhancement of synaptic and/or proteostasis function in PV interneurons prevents or delays the age-related degeneration of these cells.

## 5 Conflict of Interest

*The authors declare that the research was conducted in the absence of any commercial or financial relationships that could be construed as a potential conflict of interest*.

## 6 Author Contributions

AH: performed experiments, data analysis, design of data analysis, figure illustration, manuscript correction; CB: performed experiments, data analysis, figure illustration; assisted in writing the manuscript; DP: designed and performed experiments, data analysis, figure illustration; JD: designed and performed experiments, data analysis, figure illustration; JG: designed and performed experiments, data analysis, figure illustration; LG: performed experiments, data analysis, figure illustration; JL: performed experiments, figure illustration, manuscript correction; TP: provided help with conceptual design, discussion of results; TL: data analysis, design of data analysis; GS: performed experiments; LB: performed experiments, data analysis; JM: designed and performed experiments, data analysis, manuscript correction; MS: conceptual design, assisted in writing the manuscript; AU: designed and performed experiments, data analysis, manuscript correction; GZ: conceptual design of the study, designed and performed experiments, data analysis, figure illustration, wrote the manuscript.

## 7 Funding

This work was funded by the Deutsche Forschungsgemeinschaft (DFG, German Research Foundation) - 368482240/GRK2416 associated to GZ and MS, as well as by the DFG (ZI 1224/8-1) and the IZKF Jena, both associated to GZ; in addition to the DFG (MA-3975/2-1) associated to JM.

## 8 Acknowledgements

We thank Susanne Luthin and Fabian Ludewig from the transcriptome analysis laboratory Goettingen for excellent technical assistance. Moreover, we thank Katrin Schubert from the FACS-core facility of the FLI Jena.

## 1 Data Availability Statement

The datasets for this study can be found in the GEO database **[Series GSE145026]**.

