## Supplementary material for "DNA methyltransferase 1 (DNMT1) function is implicated in the age-related loss of cortical interneurons"

### 1 Supplementary Data

Supplementary Material should be uploaded separately on submission. Please include any supplementary data, figures and/or tables. All supplementary files are deposited to FigShare for permanent storage and receive a DOI.

Supplementary material is not typeset so please ensure that all information is clearly presented, the appropriate caption is included in the file and not in the manuscript, and that the style conforms to the rest of the article. To avoid discrepancies between the published article and the supplementary material, please do not add the title, author list, affiliations or correspondence in the supplementary files.

**Supplementary Methods**

**Cell Culture**

Neuro-2a cells (N2a) were cultured in high-glucose DMEM supplemented with 100 U/mL penicillin, 100 µg/mL streptomycin and 10% FBS at 37 °C, 95% H_2_O and 5% CO_2_.

**Transfection with siRNA Oligos**

For siRNA transfections of N2a cells, reverse lipofection with Lipofectamin© 2000 (Thermo Fisher Scientific, USA) according to the manufacturer’s protocol and as described in Zimmer et al. (2011) was applied using 15 nM control siRNA (BLOCK-iT Alexa Fluor red or green fluorescent oligo, Invitrogen, USA) and 30 nM *Dnmt1* and *Rab7* siRNA (Santa Cruz Biotechnology) for 5 h in Opti-MEM I Reduced Serum Medium without antibiotics (Thermo Fisher Scientific). 1 day prior to transfection, N2a cells were plated on coverslips coated with 19 μg/mL laminin (Sigma-Aldrich, Germany) and 5 μg/mL poly-L-lysine (Sigma-Aldrich).

**Quantitative Reverse Transcription PCR**

RNA was isolated with Trizol®Reagent (Life Technologies, USA) according to manufacturer’s guidelines and was used for cDNA synthesis using the SuperScript III. Quantitative PCR reactions for analyzing Dnmt1, Rab7 and Rps29 quantities were performed using 10 ng cDNA of each sample and the innuMix qPCR DSGreen Standard Mastermix (Jena Analytik, Germany) on the qTOWER3 G thermocycler (Jena Analytik, Germany).

The following primer sequences (indicated as 5’ → 3’) were used: *Dnmt1* forw. GGT GAG CAT CGA TGA GGA GA, rev. GCT GTG ACC CTG GCT AGA TA; *Rab7* forw. TAG GAG CGG ACT TTC TGA CC, rev. CAC CTC TGT AGA AGG CCA CA *Rps29* forw. GAA GTT CGG CCA GGG TTC C, rev. GAA GCC TAT GTC CTT CGC GT; data were normalized to Rps29. Results were analyzed via the ΔΔCt-method.

**Muscle preparation**

Mice were deeply anesthetized by intraperitoneal injection of 50% chloral hydrate in phosphate buffered saline (PBS; pH 7.4; 2.5 µg chloral hydrate per g body weight).

For hematoxylin and eosin (HE) as well as laminin staining, freshly prepared extensor digitorum longus (EDL) muscle and soleus muscle from the hind limbs were immediately frozen in embedding media (Tissue-Tek® OCT compound with 10% sucrose) in liquid nitrogen and stored at -80°C. Cryosections of frozen muscles 12 µm in thickness were cross-sectionally cut using a cryostat (Microm, Thermo Fisher Scientific, U.S.A.).

**HE-Staining, immunohistochemistry and immunocytochemistry**

For HE-staining, muscle sections were fixed in cold acetone and then stained automatically using a tissue stainer (Autostainer XL, Leica Microsystems, Germany) prior to automatic mounting in CV Mount (Leica Microsystems).

For Laminin staining, muscle sections were fixed in 2% PFA, permeabilized with 0.1%TritonX and 0.1 M glycine in PBS (pH 7.4) and blocked for 1 h with 5% horse serum in PBS (pH 7.4). Primary antibody was applied overnight at 4°C and the secondary antibody was applied for 45 min at room temperature prior to DAPI nuclear staining (1:5000 in 1xPBS; pH 7.4, Sigma-Aldrich, Germany) for 5 min. Stained muscle sections were mounted in Permafluor (Thermo Fisher Scientific).

For staining of neuromuscular junctions, freshly prepared EDL muscles were fixed in 2% PFA for 15 min and washed with PBS (pH7.4). Fiber bundles consisting of 4-5 myofibers were prepared and used for further analyses. After permeabilization in 0.1%TritonX in PBS (pH 7.4) for 10 min at room temperature, samples were blocked with 5% horse serum in PBS (pH 7.4) for 1 h followed by an incubation with bungarotoxin-Alexa 488 (1:500, Invitrogen) and primary antibodies over night at 4°C. After washing with PBS (pH7.4), single myofiber bundles were incubated with the corresponding secondary antibodies for 1 h at room temperature prior to DAPI staining (10 µg/ml, Invitrogen) for 5 min. Stained muscle fibers were washed with PBS (pH7.4) and mounted in Permafluor (Thermo Fisher Scientific).

Brain sections were labeled with antibodies according to immunohistochemistry methods in the main manuscript.

N2a cells were permeabilized with 0.2% Triton X-100 in PBS (pH7.4) for 10 min prior to blocking with 5% normal goat serum in PBS (pH7.4) for 1 h. Primary antibodies were applied overnight at 4°C, secondary antibody for 1 h at RT. After nuclei staining with DAPI (Molecular Probes) for 5 min, coverslips were embedded in Mowiol (Carl Roth). If not annotated differently, all steps were performed at room temperature.

Following primary antibodies were used: rabbit anti-Laminin, (1:1000, Abcam), mouse anti-Neurofilament (undiluted, 2H3, DSHB), mouse anti-synaptic vesicle protein 2 (undiluted, SV2, DSHB), mouse anti-NeuN (1:500, Millipore), rabbit anti-Ubiquitin (1:600, Dako).

Following secondary antibodies were applied: goat Cy5 anti-mouse (1:1000, Thermo Fisher Scientific), goat Cy5 anti-rabbit (1:1000, Thermo Fisher Scientific), goat Alexa647 anti-rabbit (1:1000, Invitrogen), goat IgG-Alexa647 anti-mouse (1:1000, Invitrogen) and donkey Alexa 488 anti-rabbit (1:500, Jackson Immunoresearch)

**Microscopy and image analysis**

Images of stained muscle and brain tissue were taken with an inverted transmittedlight microscope Axio Cellobserver Z1 equipped with MosaiX module for tile scanning and Apotome for confocal like imaging (Carl Zeiss Microscopy, Germany).

Based on images taken from the Laminin-stained muscles the diameters of 50 fibres were measured and mean fibre diameters were compared.

### 2 Supplementary Figures and Tables

**Supplementary Figure 1: Analysis of the density of NeuN-positive neurons in young and aged neocortex**

(**a**-**c**) Immunohistochemistry against the pan-neuronal marker NeuN in motor (**a**), somatosensory (**b**) and visual cortical areas (**c**) of 2 months and 16 months old mice, quantified in **d**. Scale bars: 50 µm.

**Supplementary Figure 2**

***Dnmt1* deficiency causes no obvious alterations in the cerebellum, in skeletal muscle tissue or in the muscle innervation**

(**a**, **b**) *In situ* hybridization of sagittal sections of the cerebellum from C57BL/6 mice (3 months) using riboprobes directed against *Dnmt1* (**a**) and *Pvalb* (**b**) transcripts revealed expression of both transcripts in the Purkinje cell layer of the cerebellum. (**c**) Schematic illustration of the location of microphotographs shown in (**d**, **e**) in the cerebellum of sagittal brain sections. (**d**, **e**) The analysis of Purkinje cells labeled by tdTomato in *Pvalb*-Cre/tdTomato/*Dnmt1* wild-type (WT) and knockout (KO) mice counted in the cerebellum of sagittal brain sections of young (6 months; n=3 mice per genotype) and aged (16 months, n=3 mice per genotype) mice showed no differences in cell number (**e**) (**f**, **g**) Representative images of muscle cross-sections of the extensor digitorum longus (EDL) muscle (**f**) and the soleus muscle (**g**) of 6 and 16 months old *Pvalb*-Cre/tdTomato/*Dnmt1* wild-type (WT) and knockout (KO) mice. Laminin (red) / DAPI (blue) staining showed no difference in mean fiber diameter quantified in (**h**) between WT and KO (6 months: n = 3 mice per genotype, 16 months: n = 4 mice for WT, 3 mice for KO). (**i**, **j**) Representative images of hematoxylin and eosin-stained muscle cross-sections of the EDL muscle (**i**) and the soleus muscle (**h**) of 6 and 16 months old *Dnmt1* WT and KO mice revealed no obvious muscle damage (6 months: n = 3 mice per genotype, 16 months: n = 4 mice for WT, 3 mice for KO). (**k**) Representative images of neuromuscular junctions from EDL muscle of 6 and 16 months old *Dnmt1* WT and KO mice stained for nicotinic acetylcholine receptors (with bungarotoxin), neurofilaments (NF) and synaptic vesicle protein 2 (SV2) show the proper innervation of the neuromuscular junctions of WT and KO mice. (6 months: n = 3 mice per genotype, 16 months: n = 4 mice for WT, 3 mice for KO). Scale bars: 250 µm in (**a**, **b**); 50 µm in (**d**), 100 µm in (**f**, **g**, **i**, **j**); 10 µm in (**k**)

**Supplementary Figure 3**

**Transcriptome and methylation analysis of FACS-enriched adult and aged *Dnmt1*-deficient and wild-type *Pvalb*-expressing cortical interneurons.**

(a) FACS-events determined per cortical hemisphere for young and old *Pvalb*-Cre/*tdTomato* wild-type mice. (**b-d**) Gene expression analysis of young (6 months) and old (18 months) FACS-enriched *Pvalb*-Cre/*tdTomato* wild-type (WT) and *Pvalb*-Cre/*tdTomato* *Dnmt1 loxP^2^* (knockout, KO) cortical interneurons was performed by RNA-sequencing (pooled samples from N = 6 WT and KO mice at 6 months; and N = 9 WT and N = 12 KO mice at 18 months were analyzed in technical duplicates). (**b**-**d**) Principal component analysis (PCA) illustrates the segregation of the different samples in the dendrogram (**b**) and the PCA plot (**d**), whereby the first PC is determined by more than 80% of the variance (**c**).

**Supplementary Figure 4**

**DNMT1 affects endocytic-based degradation**

(**A**) Verification of *Dnmt1* and *Rab7* siRNA efficiency in N2a cells measuring the expression levels by quantitative RT-PCR after 24 h of siRNA or control siRNA treatment (normalized to *Rps29*) and *ActB* expression). (**b**-**d**) CD63 (magenta) and LAMP1 (green) antibody staining in CB cells (**b**) and N2a cells (**c**) after control and *Dnmt1* siRNA treatment for 1 div. The white rectangles illustrate the locations of the magnified parts of the processes. Quantification of fluorescence intensities is shown in (**d**) for all cell types. (**e**, **f**) Ubiquitin immunostaining in the motor cortex of sagittal brain sections of 6 months old *Pvalb*-Cre/tdTomato *Dnmt1* wild-type (WT) and knockout mice (KO; tdTomato in red, Ubiquitin in green). dtTom = tdTomato, Ub = Ubiquitin. Scale bars: 20 µm in (**b, e, f**); 10 µm in (**c**).

**Legend for supplementary tables and movies:**

**Supplementary Movies 1a, b and 2a, b:**

**DNMT1 influences endosomal transportation in cerebellar granular (CB) cells.**

Live cell imaging of CB cells with a CD63-GFP overexpression revealed a significantly faster rate of retrograde transport in the DNMT1 knockdown cells (Movie 1a, b) compared to control cells (Movie 2a, b). Scale bar 10 µm. CB cells were imaged every 5 min for 20 mins, video is produced with 3 frames per sec.

#### Supplementary Tables:

**Supplementary Table S1 (Table S1_DEG aging in WT):**

**Differential gene expression analysis between young and old *Pvalb*-*Cre/tdTomato* wild-type interneurons**.

RNA sequencing of 6 months old and 18 months old Pvalb-Cre/tdTomato wild-type interneurons of the cerebral cortex was performed. Differentially expressed genes (DEG) are listed in sheet 1-1. Gene ontology (GO) analysis was performed with DAVID. Sheet 1-2 provides an overview of GO-terms found for upregulated genes in aged wild-type interneurons, while in sheet 1-3 collects the GO-terms for down-regulated genes in aged wild-type interneurons.

**Supplementary Table S2 (Table S1_DEG aging in KO):**

**Differential gene expression analysis between young and old *Pvalb*-*Cre/tdTomato/Dnmt1 loxP^2^* interneurons**.

RNA sequencing of 6 months old and 18 months old *Pvalb-Cre/tdTomato/Dnmt1 loxP^2^* interneurons of the cerebral cortex was performed. Differentially expressed genes (DEG; Benjamini adjusted p<0.05; ⎜log2fc ⎜>1) are listed in sheet 2-1. Gene ontology (GO) analysis was performed with DAVID. Sheet 2-2 provides an overview of GO-terms found down-regulated genes in aged *Dnmt1*- deficient interneurons. As only very few genes were found up-regulated upon aging in

*Pvalb-Cre/tdTomato/Dnmt1 loxP^2^* interneurons, GO-analysis was not applicable.

**Supplementary Table S3 (Table S3 DEG DMG OL yng WT old WT_DEG with GO):**

**Analysis of genes which were changed in gene expression and DNA methylation levels** between tdTomato-positive interneurons of the cerebral cortex of 6 months old and 18 months old *Pvalb-Cre/tdTomato* wild-type mice as revealed by RNA sequencing and MeDIP sequencing. Sheet 3-1 provides an overview of genes which were significantly changed in expression **(**Benjamini adjusted p<0.05) and DNA methylation level (adjusted p<0.05) between young and aged wild-type interneurons. Gene ontology (GO) analysis performed with DAVID is collected in sheet 3_2. Sheet 3_3 depicts the unmapped genes.

**Supplementary Table S4 (Table S4 DEG DMG OL Old WT Old KO DEG with GO):**

**Analysis of genes which were changed in gene expression and DNA methylation levels** between tdTomato-positive interneurons of the cerebral cortex of aged (18 months) old *Pvalb-Cre/tdTomato* wild-type and *Pvalb-Cre/tdTomato/Dnmt1 loxP^2^* knockout mice as revealed by RNA sequencing and MeDIP sequencing. Sheet 4-1 provides an overview of genes which were significantly changed in expression **(**Benjamini adjusted p<0.05) and DNA methylation level (adjusted p<0.05) between aged wild-type and *Dnmt1*-knockout interneurons. Gene ontology (GO) analysis performed with DAVID is collected in sheet 4_2. Sheet 4_3 depicts the unmapped genes.

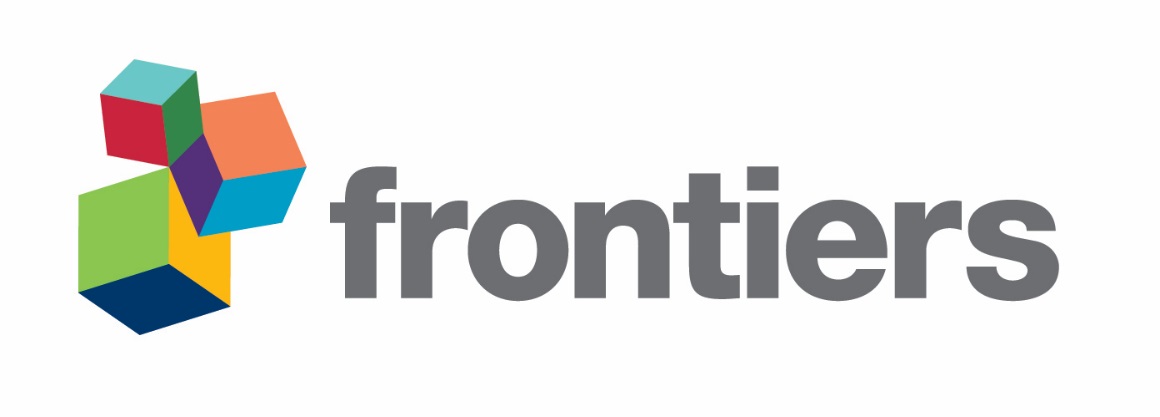
