## Supplementary figures and images for "DNA methyltransferase 1 (DNMT1) function is implicated in the age-related loss of cortical interneurons"

### Supplementary Figure 1

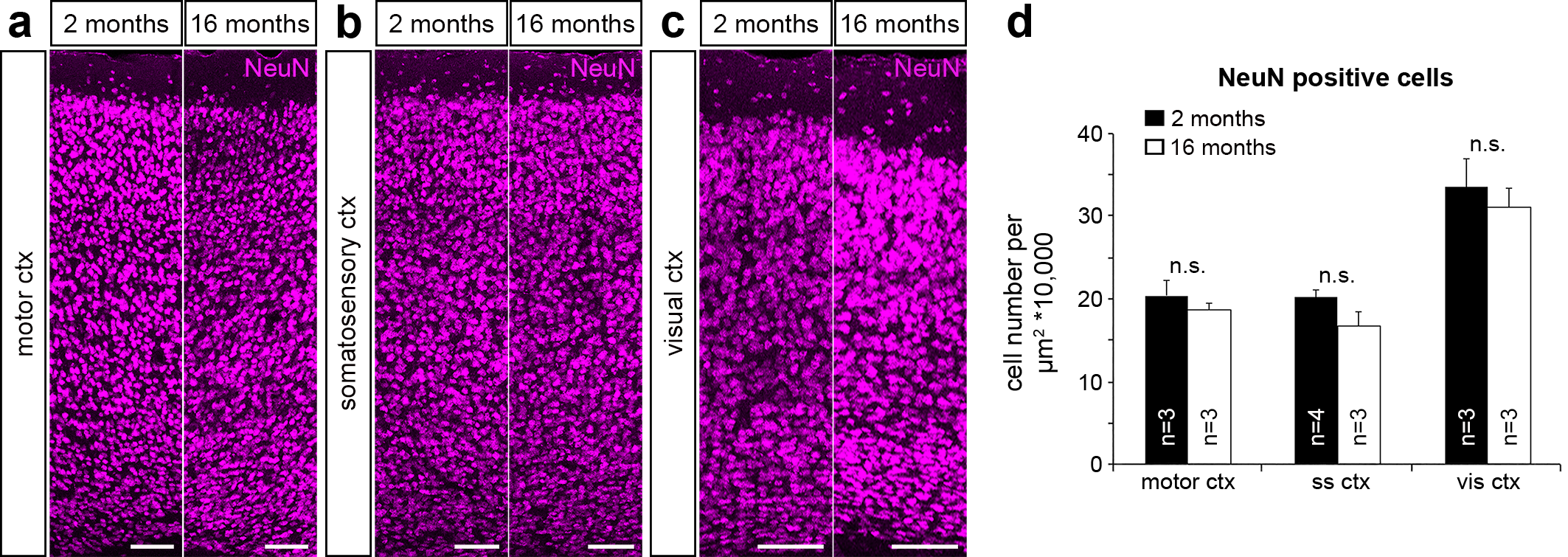

### Supplementary Figure 2

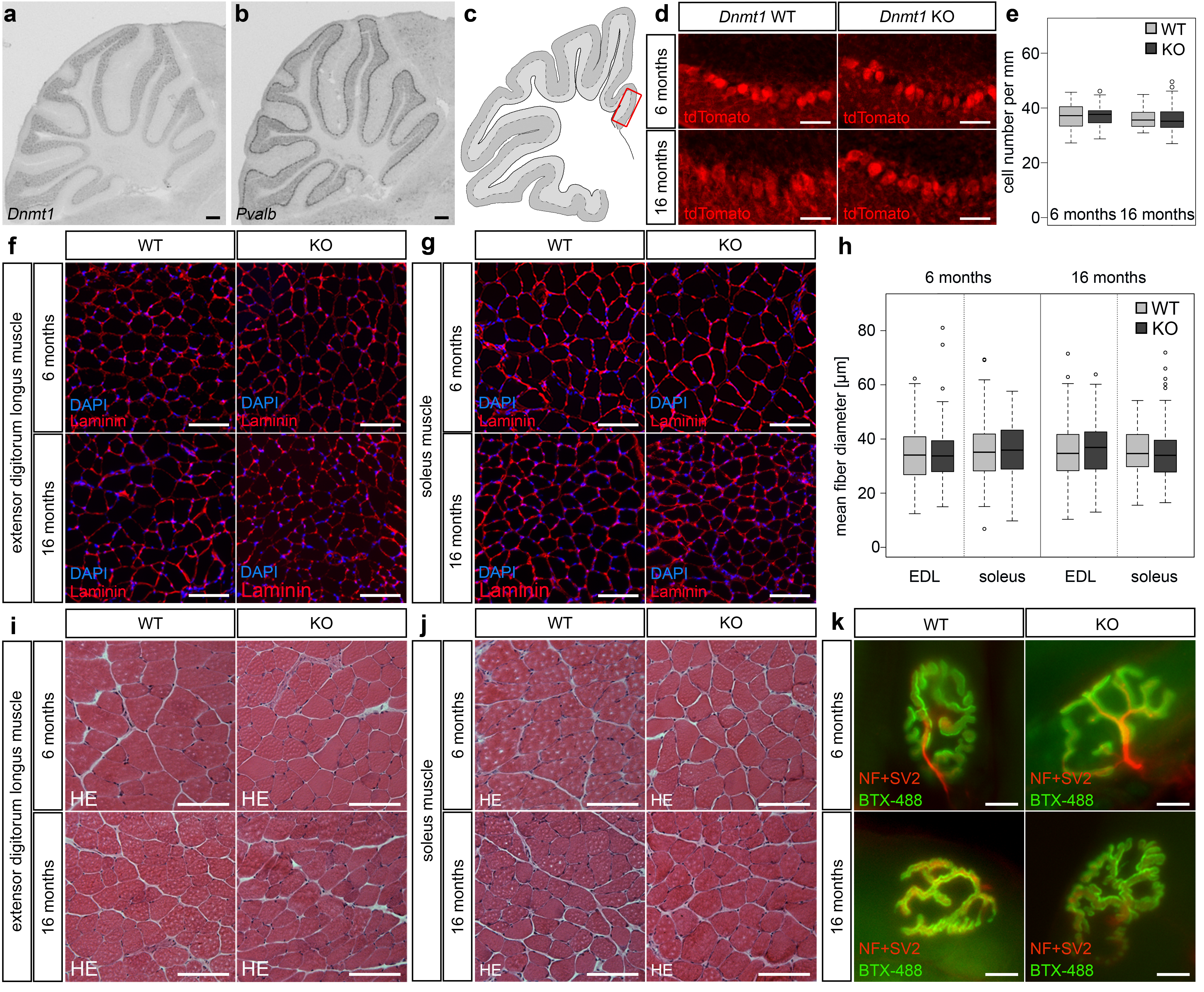

### Supplementary Figure 3

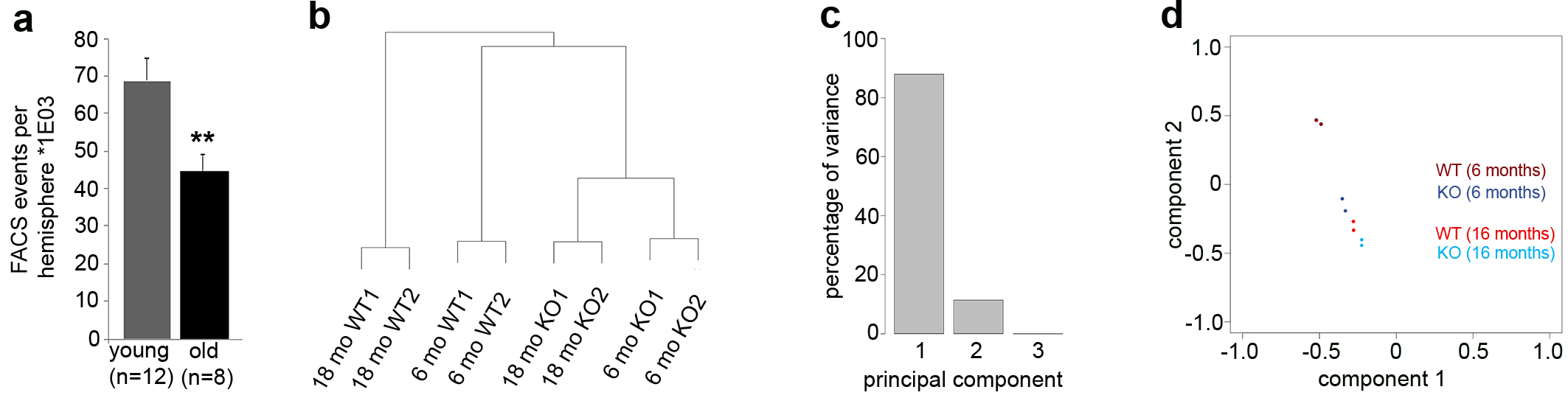

### Supplementary Figure 4

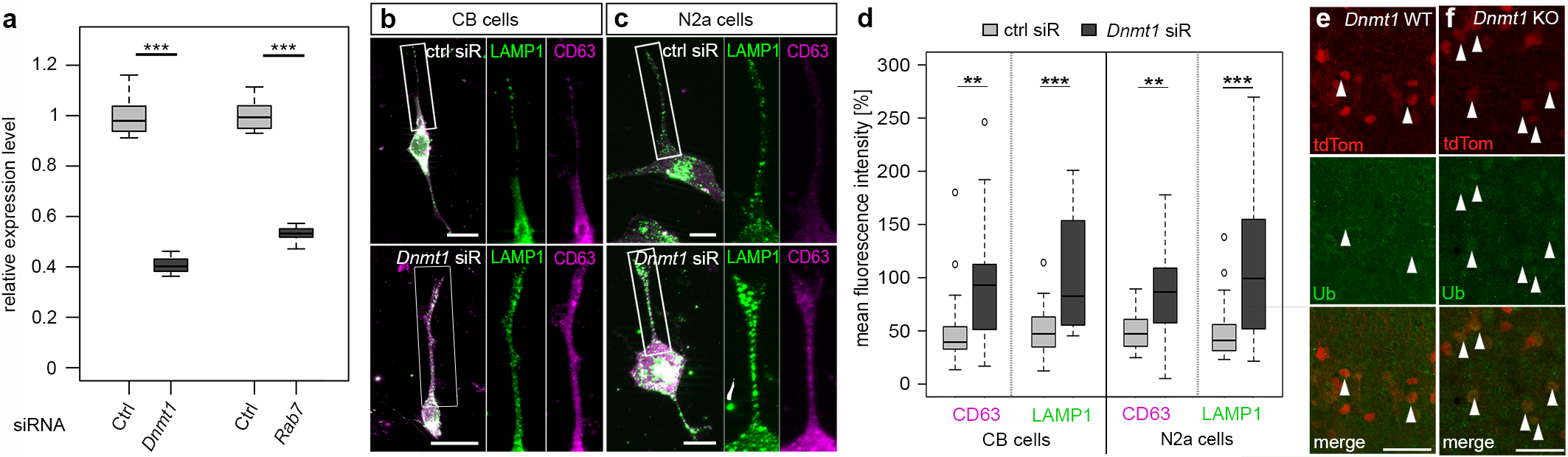
